# Precise engineering of chimeric antigen receptor expression levels defines T cell identity and function

**DOI:** 10.1101/2025.02.10.637482

**Authors:** Andrew S. Ramos, Sylvain Simon, Jesus Siller-Farfan, Anusha Rajan, Santiago Revale, Elena Zanchini di Castiglionchio, Oana Pelea, Tudor A. Fulga, Stanley Riddell, Omer Dushek, Yale S. Michaels, Tatjana Sauka-Spengler

## Abstract

Chimeric Antigen Receptor (CAR) T therapy is a potent treatment for haematological malignancies, but T cell exhaustion reduces its efficacy in many patients. Although high CAR transgene levels appear to drive T cell exhaustion, the relationship between CAR expression levels, T cell function, and transcriptional identity is yet to be mapped at high resolution. Here, we harness a high-resolution microRNA-based control system to precisely modulate CAR transgene expression levels and assess the impact on T cell activation, gene expression and function. By post-transcriptionally modulating CAR abundance, we show that differential CAR levels significantly impact T cell proliferation, cytokine production and tonic signalling. T cells with high CAR expression become strongly activated even at low target antigen densities, while those with low CAR expression are triggered only by high concentrations of their target. Single-cell RNA sequencing of primary T cells expressing a broad range of CAR transcript levels revealed global transcriptional programmes that become dysfunctional with increased CAR abundance, expanding our understanding of T cell exhaustion. Notably, we identified a narrow CAR expression range where the exhaustion transcriptional state is not triggered, demonstrating that T cell exhaustion can be controlled by fine-tuning CAR levels. This work demonstrates that CAR expression levels are key determinants of T cell transcriptional identity and function and introduces a tractable method to precisely tune CAR expression and T cell activity.

## 1 Introduction

Chimeric antigen receptor (CAR) T cells have emerged as a potent cancer treatment, achieving high remission rates for haematological disorders and leading to the FDA approval of six CAR-T products [1, 2, 120–122]. However, several limitations hamper their widespread translation into clinical benefit for cancer patients. Notably, a large proportion of patients receiving treatment experience toxic side effects, such as cytokine release syndrome (CRS) and neurotoxicity, whose exact pathophysiologies are not fully understood but are suggested to be caused by excessive or chronic activation of CAR T cells [3, 4, 85, 87, 118–120, 123–126]. While the majority of toxicities are low grade and have established management protocols, there is a fraction of patients in which side effects are life-threatening [85, 118, 120]. Importantly, managing such symptoms is expensive and extends median hospital stay [128–133], highlighting the critical need to prevent such side effects altogether. Furthermore, T cell exhaustion reduces the success of CAR T cell therapy, affecting both the response rates in haematological cancers where CAR Ts have become standard treatment and their application to solid cancers [5, 116, 117]. CAR T cell therapy aims to redirect T cells towards a specific antigen expressed on tumour cells by enforcing the expression of a chimeric antigen receptor synthetic protein. Manufacturing protocols for FDA-approved CAR T therapies include use of constitutive promoters (e.g., CMV [6], EF1*α* [108, 109], MND [6, 9, 109], etc.) coupled with retroviral transduction methods (e.g., gamma retroviral or lentiviral) to integrate the CAR transgene into the T cell genome. This results in the supraphysiological expression of CAR proteins on the cell surface compared to endogenously expressed T cell receptors [6–11]. High CAR density has been linked with tonic signalling due to CAR-scFv self-association [7, 12, 13], which is suggested as a driver of T cell exhaustion [6, 8, 10, 12, 13].

A growing number of studies have shown that CAR expression levels play a key role in T cell phenotype, function, transcriptional identity, and epigenetic landscape [8, 10, 27, 133]. Indeed, CAR-Ts produced by introducing a CD19 CAR transgene under the control of the endogenous TRAC locus result in lower and more consistent CAR expression levels [8]. These CAR-T cells exhibit enhanced *in vivo* anti-tumour potency, reduced differentiation, lower expression of inhibitory receptors PD-1, CTLA-4, LAG3, and reduced T cell exhaustion compared to CAR-Ts produced using standard retroviral manufacturing. Another study found that retrovirally manufactured CAR-Ts sorted into low and high CAR expression populations displayed stronger gene expression signature levels for tonic signalling, exhaustion, and activation in high CAR expressing T cells [10]. These studies, while indicative, are limited by assessing only two levels of CAR expression at low resolution, highlighting the need to understand the precise relationship between CAR expression level and T cell behaviour and state. We hypothesised that by finely tuning CAR expression to precise, incremental levels, we could quantitatively analyse the relationship between CAR expression and T cell states.

To test this hypothesis, we used MicroRNA silencing-mediated fine-tuners (miSFITs) to tune CAR expression levels in T cells [14–16]. MicroRNAs (miRNA) are endogenous, short non-coding RNAs that regulate gene expression post-transcriptionally by recruiting RNA-induced silencing complex (RISC) to cellular RNAs containing complementary miRNA response elements (MREs). The magnitude of repression depends on the degree of complementarity between a miRNA and its target MRE. miSFITs have emerged as a robust technology that harnesses microRNAs to precisely control gene expression levels [14–16]. miSFITs are artificial MREs in which point mutations have been introduced into the sequences, changing the degree of complementarity between miRNA and MRE, thus altering their repressive strength and modulating gene expression levels in a predictable manner [14–16].

In this study, we harnessed miSFIT technology to finely tune CAR expression, enabling a comprehensive, high-resolution analysis of the relationship between CAR abundance and T cell function and transcriptional state. We used a library of miSFIT lentiviral vectors to achieve precise, stepwise expression control of a NY-ESO-1-targeting CAR in immortalised cell lines and primary T cells. Through *in vitro* co-cultures, we investigated the role of CAR expression in T cell activation, demonstrating that tuned CAR expression dictates both the magnitude of activation and the proportion of activated T cells. We further show that CAR expression levels modulate the antigen density threshold required for T cell activation. Additionally, we reveal that CAR expression plays a defining role in T cell function, as evidenced by direct correlations between CAR expression level and T cell proliferation, cytokine production, killing capacity, and tonic signalling. Using single cell RNA-sequencing, we identify fundamental biological processes that become progressively dysfunctional with increasing CAR expression levels, including RNA processing, cell proliferation, and metabolic function, expanding our understanding of T cell exhaustion. This study highlights the importance of CAR expression levels as a key determinant of CAR-T cell function and identity, and may inform future efforts to modulate CAR T therapy through precise expression modulation.

## 2 Results

### 2.1 miSFITs precisely tune CAR expression levels at high resolution

To begin fine-tuning CAR expression, we adapted a previously validated miR-17 responsive miSFIT library [14]. miR-17 is an attractive target as it is a broadly expressed miRNA, including in CD8+ T cells [17, 18]. We used a miSFIT library incorporating 15 single and di-nucleotide miR-17 target site variants (**Figure 1a**), which have been shown to robustly tune gene expression levels [14–16]. To achieve a broad dynamic range, we also included single, duplicate, and quadruplicate perfectly complementary miR-17 miSFITs, referred to as 1x, 2x, and 4x Perfect, respectively, which we expected to strongly repress CAR expression. We also included a variant with minimal complementarity to miR-17 (Scramble miSFIT), expected to result in the highest, unregulated level of CAR expression, serving as a control comparable to the unsilenced CAR-T vectors currently in clinical use.

**Fig. 1:**
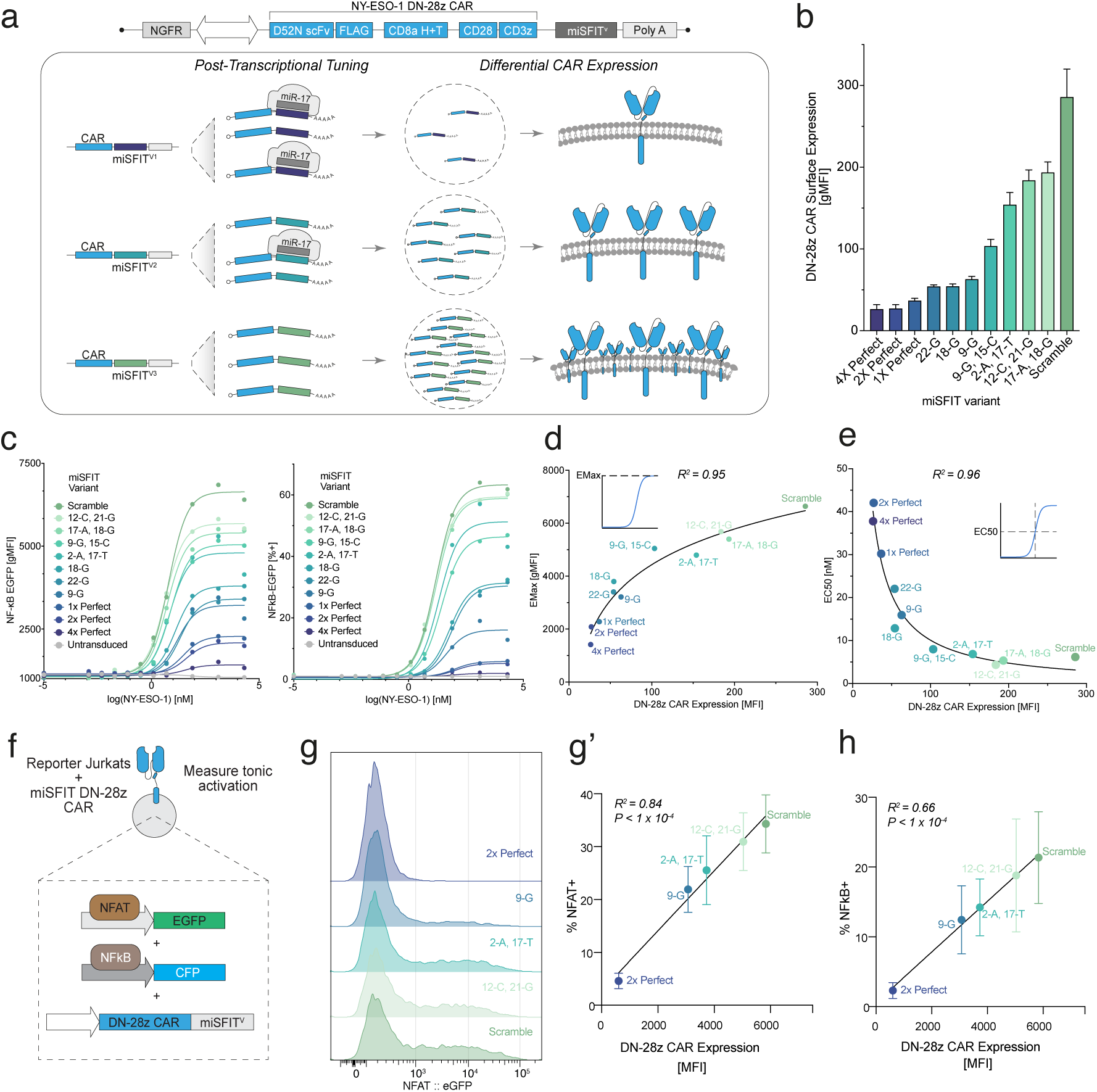
miSFITs robustly and precisely tune DN-28z CAR expression in Jurkat T cells. **a)** DN-28z CAR-miSFIT expression library used to tune CAR expression. A miSFIT (a target site for the endogenously expressed miR-17, containing 0, 1 or 2 mismatches) is introduced into the 3’ UTR of each vector,targeting the DN-28z CAR transcript for post-transcriptional regulation by endogenous miR-17. **b)** Average expression of each DN-28z CAR miSFIT cell line. N=3 biological replicates, error bars depict standard deviation. **c)** DN-28z CAR miSFIT cell lines were co-cultured with synthetic target cells, pulsed with serial dilutions of NY-ESO-1 peptide. Each DN-28z CAR miSFIT cell line contained a GFP reporter of NF-*κ*B activity (**Extended Data** Figure 3). **c)** Hill model fitting of EGFP average MFI and %EGFP+ cells for each miSFIT cell line. Each colour represents a different miSFIT cell line. **d)** Logarithmic regression analysis between average CAR expression and EGFP MFI EMax. **e)** Inverse regression analysis between average CAR expression and EGFP MFI EC50. **f-h)** To evaluate the role of CAR expression on tonic signalling, we transduced a panel of DN-28z CAR miSFITs into a Jurkat cell line containing a reporter of NF-*κ*B and NFAT and measured basal reporter activity by flow cytometry. **f)** Schematic of experimental setup to assay tonic signalling. **g)** Representative flow cytometry histograms of basal NFAT reporter activity. **g’ & h)** Linear regression analysis between %NF-*κ*B+ and %NFAT+ cells and CAR expression of CAR-miSFIT Jurkat cell lines. R^2^ refer to the goodness of model fit.

We cloned miSFIT sequences downstream of a New York squamous cell carcinoma 1 antigen (NY-ESO-1) targeting CAR. NY-ESO-1 is a cancer-testis antigen and a promising target in a range of solid tumours (e.g., ovarian cancer and metastatic melanoma) [19]. We selected the reported DN-28z NY-ESO-1 targeting second-generation CAR, given its enhanced specificity, antigen binding and micromolar affinity comparable to endogenous T cell receptors (TCR) [20]. We have extensively studied the DN-28z CAR in numerous CAR formats and compared directly to the TCR [94]. Further, the DN-28z CAR is a peptide-major histocompatibility complex (pMHC) targeting CAR, enabling antigen titrations, which is not readily available for surface antigens like CD19. Additionally, we incorporated a FLAG epitope tag to facilitate the quantification of cell surface CAR abundance in subsequent experiments.

We cloned the DN-28z CAR into a bi-cistronic expression vector library that encodes both a reporter gene, nerve growth factor receptor (NGFR), and a separate transcription unit where the CAR is placed under post-transcriptional control of one of the miSFIT variants (**Figure 1a**).

To test whether miSFIT variants could tune DN-28z CAR expression, we performed a pooled transduction experiment in Jurkat T cells, an immortalised cell line used in T cell activation studies [21] we validated to expression miR-17 (**Extended Data** Figure 1a). 72 hours post-transduction, we extracted the genomic DNA (gDNA) and mRNA and subjected them to targeted, high-throughput sequencing. To calculate DN-28z transcript expression for each miSFIT, we normalised the count of each variant sequence in the mRNA pool by its respective count in gDNA pool, which achieved a stepwise modulation of DN-28z CAR transcript expression levels (**Extended Data** Figure 1b). miSFITs provided clear, incremental control of CAR transcript levels with a dynamic range of 0.43-1.96 normalised transcript counts with an average interval of 0.085.

To verify tuning at the protein level, we generated 11 polyclonal cell lines, each expressing a different CAR-miSFIT vector (**Extended Data** Figure 2). Flow cytometry confirmed that miSFITs exerted a stepwise control of expression levels with high reproducibility (**Figure 1b** and **Extended Data** Figure 2b). The miSFIT panel provided an 11-fold dynamic range between the highest and lowest level of DN-28z CAR expression, with 1X, 2X, and 4X perfectly complementary MREs yielding the lowest levels of CAR expression and intermediate miSFIT variants conferring incremental increases. Collectively, both analyses at transcript and protein levels demonstrate that miSFIT approach enables robust and precise tuning of CAR expression. We next focused our efforts on understanding the role of CAR expression levels in T cell response. We used a Jurkat cell line with an NF-*κ*B-reporter to measure the impact of CAR expression on T cell activation [20]. NF-*κ*B is a family of transcription factors whose signalling cascades have multiple roles in T cell activation, such as the upregulation of early T-cell activation markers [22, 23].

To simulate T cell activation mediated by antigen engagement, we co-cultured DN-28z CAR-miSFIT Jurkats with an artificial antigen presentation system. This system is based on the Chinese Hamster Ovary-K1 (CHO-K1) [24] cell line engineered to express human CD58 (hCD58, which strengthens the interaction between CAR-T and target cell by binding to T cell surface antigen CD2 [25, 26]) and human leukocyte antigen A*02 (HLA-A*02). We pulsed CHO-K1 cells with NY-ESO-1_157_*_−_*_165_ peptide, which is recognised by the DN-28z CAR [20]. Subsequently, we co-cultured DN-28z CAR-miSFIT cell lines with CHO-K1 cells transduced with various concentrations of target peptide and measured T cell activation by flow cytometry.

T cell activation profiles displayed a very strong and clear association with CAR expression levels (**Extended Data** Figure 3). Fitting a Hill model to the NF-*κ*B reporter activity to CAR expression revealed distinct, tuned levels of expression with the different CAR-miSFIT Jurkat cell lines (**Figure 1c**, **Extended Data** Figure 3). Notably, the magnitude of activation (MFI of EGFP reporter) and the proportion of cells activated (%EGFP+) both depended strongly on CAR expression levels (**Figure 1c**). This is notable, as previous evidence suggests that TCR-mediated T cell activation is a binary process, whereby the proportion of T cells that are activated changes in response to target antigen concentration, but the magnitude of the activation state is fixed [29]. Yet, our finding that CAR expression impacts the magnitude of NF-*κ*B signalling in a continuous, rather than binary manner, is consistent with a previous report [27]. The relationship between the magnitude of activation (EMax of MFI) and CAR expression closely followed a logarithmic model R^2^ = 0.95 (**Figure 1d**). This implies that tuning CAR expression is an excellent strategy to control the magnitude of activation. We detected a very strong inverse relationship between EC50 (the concentration of antigen required to elicit half-maximal T cell activation) and CAR expression levels (R^2^ = 0.96, **Figure 1e**), further reinforcing the notion that tuning CAR expression may represent a promising strategy to control the magnitude of activation upon exposure to varying levels of target antigen expression.

### 2.2 Tuned CAR expression controls T cell functions

High levels of CAR expression promote tonic signalling which impacts T cell function and eventually leads to T cell exhaustion [6, 8, 10, 12, 13, 33]. However, the dose-response relationship between CAR expression levels and tonic signalling has not been mapped. To address this gap, we utilised an engineered Jurkat T cell line harbouring reporters of NF-*κ*B and NFAT transcription factors activity as a readout of tonic signalling [92]. For a panel of DN-28z CAR-miSFITs, we measured basal reporter activity in the absence of target antigen by flow cytometry and observed a significant correlation with CAR expression (**Figure 1f-h**, **Extended Data** Figure 4). We observed a strong linear relationship between both %NF-*κ*B+ and %NFAT+ and CAR expression (NFAT: R^2^ = 0.84, P *<* 1 x 10*^−^*^4^ and NF-*κ*B: R^2^ = 0.66, P *<* 1 x 10*^−^*^4^, **Figure 1g’ & h**). These findings suggest that CAR expression level is a key determinant of basal T cell activation in the absence of target antigen.

Motivated by our findings using immortalised reporter cell lines, we used the miSFIT system to fine-tune CAR expression levels in primary T cells (**Extended Data** Figure 5). Initial quality controls following transduction and expansion shows precise, step-wise levels of CAR expression for different miSFIT variants in the library (**Extended Data** Figure 5).

We sorted primary T cells transduced with each miSFIT-CAR vector based on the expression of an NGFR reporter, whose expression level is not controlled by the miSFIT. We then co-cultured each T cell population with NY-ESO-1-presenting CHO-K1 target cells and measured activation by flow cytometry (**Extended Data** Figure 6). Both the magnitude and the frequency of CD69 expression (an early activation marker [34, 35]) displayed a clear dependence on CAR expression levels (**Extended Data** Figure 6), confirming that CAR expression controls T cell activation in primary T cells.

To measure the impact of CAR expression on the antigen-dose response of T cell signalling and cytokine production, we exposed a panel of DN-28z CAR miSFIT T cells to titrated concentrations of NY-ESO-1 and measured phosphorylated extracellular signal-regulated kinase (pERK) levels, **(Figure 2a & b and Extended Data** Figure 7a) and Interferon-*γ* (IFN-*γ*) production by intracellular cytokine staining (**Figure 2a & c** and **Extended Data** Figure 7b). We then fitted both pERK and IFN-*γ* measurements using a Hill-model (**Figure 2b & c, left**). pERK and IFN-*γ* activity levels varied with CAR expression (**Figure 2b, left, & c, left**). We identified a direct, inverse relationship between pERK EC50 and CAR expression (R^2^ = 0.98, **Figure 2b, middle**). However, we did not identify a strong relationship between IFN-*γ* and CAR expression with linear regression (R^2^ = 0.15, **Figure 2c, middle**). We next fitted the EMax of %pERK+ and %IFN-*γ*+ with CAR expression using linear and logarithmic regression, respectively, and found very strong relationships for both parameters (**Figure 2b, right**, pERK: R^2^ = 0.92, **Figure 2c, right**, IFN-*γ*+: R^2^ = 0.91). These data demonstrate that CAR expression modulates T cell activation across a range of antigen concentrations (**Figure 2b & c** and **Extended Data** Figure 7).

**Fig. 2:**
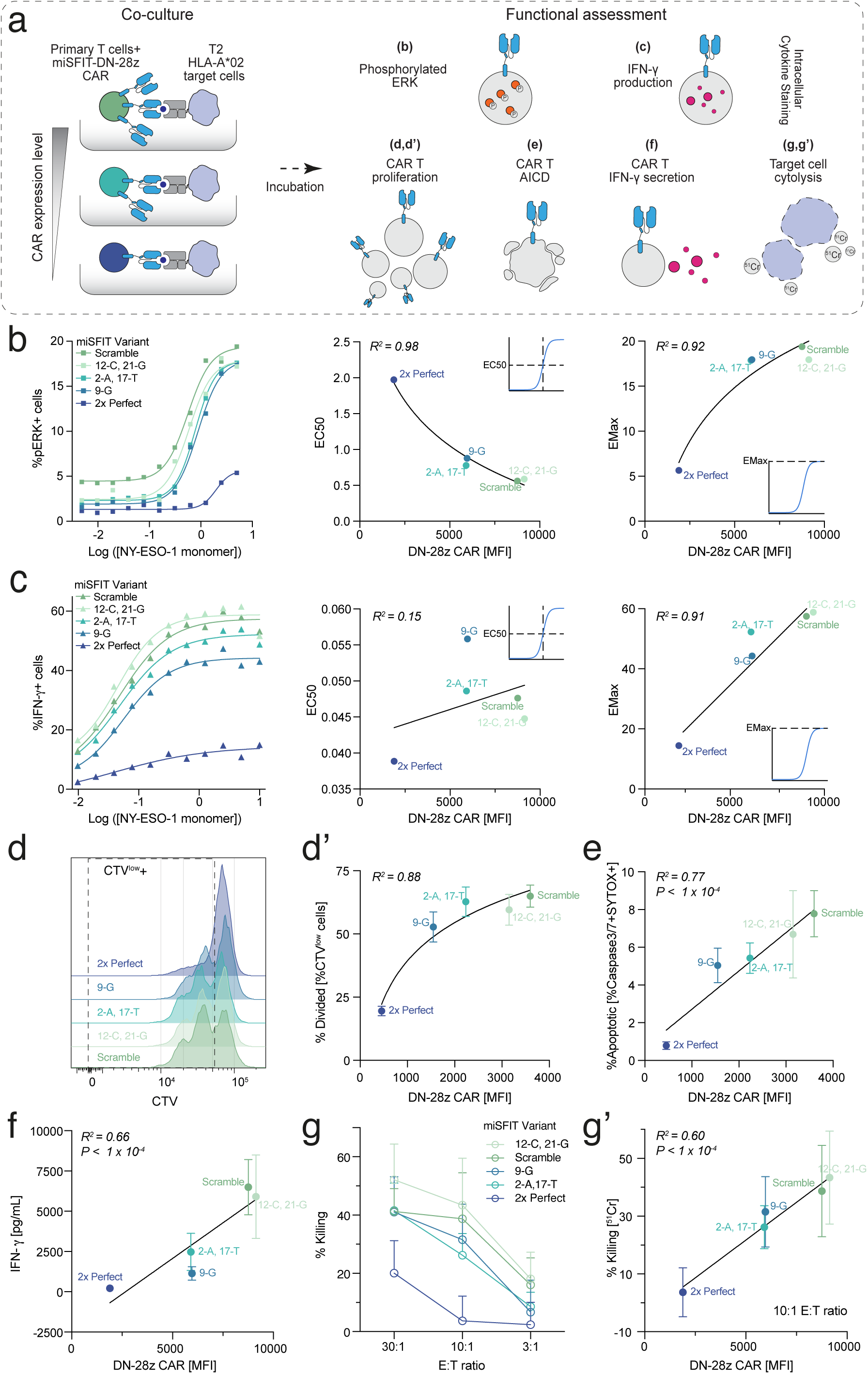
CAR expression levels are a key determinant of T cell function. **a)** Overview and experimental setup of assayed CAR T functional properties **(b-g’)**. **b & c)** We measured the impact of CAR expression on the sensitivity of T cell signalling and cytokine production by exposing a panel of DN-28z CAR miSFIT T cells to titrated concentrations of NY-ESO-1 peptide and assayed phosphorylation of extracellular signal-regulated kinases (pERK) and Interferon-*γ* production by flow cytometry. Left: Hill model fitting of %pERK+ **(b)** and %IFN-*γ*+ **(c)** for each miSFIT cell line. Centre: Regression analysis between average CAR expression of CAR-miSFIT T cells and EC50 of %pERK+ (inverse model) and %IFN-*γ*+ (linear model). Right: Model fitting between average CAR expression and EMax of %pERK+ (logarithmic model) and %IFN-*γ*+ (linear model). **d-g’)** We cocultured a panel of DN-28z CAR miSFIT T cells with T2 target cells, pulsed with 10*µ*M of NY-ESO-1 9V peptide, to assay CAR expression’s influence on T cell proliferation, activation induced-cell death, cytokine production, and cytotoxicity. Regression analysis between CAR expression and proliferation rate (**d’**, logarithmic regression), activation induced cell-death (**e**, linear), cytokine production (**f**, linear), and cytotoxicity (**g’**, linear). **d) g)** Representative flow cytometry histograms of cell proliferation as measured by CellTrace Violet dilution. **g)** Average cytotoxicity (% Killing by ^51^Cr release) per CAR-miSFIT T cell line by effector:target ratio. Error bars report standard deviation. R^2^ and P refer to the goodness of model fit parameters.

Next, we measured the impact of CAR expression levels on T cell proliferation. We co-cultured a panel of DN-28z CAR miSFIT T cells with NY-ESO-1 presenting-T2 target cells and measured T cell proliferation using CellTrace Violet (CTV) (**Figure 2a, d, & d’**, **Extended Data** Figure 8a). Proliferation increased with higher levels of CAR expression, as confirmed by logarithmic regression between % Divided and CAR expression (R^2^ = 0.88, **Figure 2d & d’**, **Extended Data** Figure 8a). Taken altogether, these results demonstrate that incremental tuning of CAR expression determines events (e.g., activation signalling cascades, cytokine secretion, proliferation) of T cell activation.

Activation-induced cell death (AICD) is a regulatory process that occurs in T cells in response to excessive activation and proliferation signals [36, 111]. As we observed a direct impact on T cell activation and proliferation rate, we examined the relationship between CAR expression levels and induction of T cell apoptosis, a functional readout of AICD. We co-cultured a panel of DN-28z CAR miSFIT T cells with NY-ESO-1 presenting T2 cells for 24 hours and measured apoptosis within the T cell population via flow cytometry (**Figure 2a & e**, **Extended Data** Figure 8b). We found a strong and significant linear relationship between CAR expression and apoptotic T cells (**Figure 2e**, R^2^ = 0.77, P *<* 1 x 10*^−^*^4^), demonstrating the direct effect of increased CAR expression on AICD.

We next investigated whether fine-tuning CAR expression levels could also fine-tune T cell effector functionality *in vitro*. To this end, we co-cultured DN-28z CAR miSFIT T cells with NY-ESO-1 presenting target cells and quantified IFN-*γ* release (**Figure 2a & f** and **Extended Data** Figure 9a). We found that IFN-*γ* concentration increased directly with CAR expression level (**Figure 2f**, R^2^ = 0.66, P*<* 1 x 10*^−^*^4^). To assess T cell cytotoxicity, we once again co-cultured our DN-28z CAR miSFIT T cells with T2 target cells at various effector-to-target (E:T) ratios and measured chromium (^51^Cr) release by target cell cytolysis (**Figure 2a, g, & g’** and **Extended Data** Figure 9b **& c**). Higher levels of CAR expression induced by the corresponding miSFIT promoted increased cell death, whereas T cells with low CAR expression levels demonstrated a notable reduction in % killing at lower E:T ratios (**Figure 2g & Extended Data** Figure 9b). This demonstrates a minimum CAR expression threshold for effective anti-tumour function. Linear regression between CAR expression and % killing confirmed the direct role of CAR expression levels in cytotoxicity (**Figure 2g’**, R^2^ = 0.60, P*<* 1 x 10*^−^*^4^). Taken altogether, these results demonstrate the pivotal influence of CAR expression on T cell effector function and suggest a potential strategy to fine-tune anti-tumour efficacy.

### 2.3 Dissecting the impact of CAR expression on the T cell transcriptome

Our data collectively reveal that CAR expression levels control T cell activation and function in a continuous, rather than binary, manner (**Figure 2**). To understand the gene expression programmes that underlie the relationship between CAR expression and T cell function, we designed a pooled, single cell transcriptional profiling experiment (**Figure 3a**). We isolated and transduced CD8+ T cells from three human donors using a pool of lentivirus containing 19 DN-28z-CAR-miSFIT variants (**Methods and Materials**, **Extended Data** Figure 10). We enriched transduced cells using magnetic-activated cell sorting (MACS) and then spiked untransduced control cells back at a defined ratio (**Methods and Materials**). We then co-cultured the transduced T cell pool for 24 hours with the CHO-K1-based activation system, recovered the CD45+ T cells from the co-cultures by FACS, and captured cells for single cell RNA sequencing (scRNA-seq). To minimise inherent technical bias, we pooled samples from three T cell donors into a single 10X channel (**Figure 3a, Methods and Materials**).

**Fig. 3:**
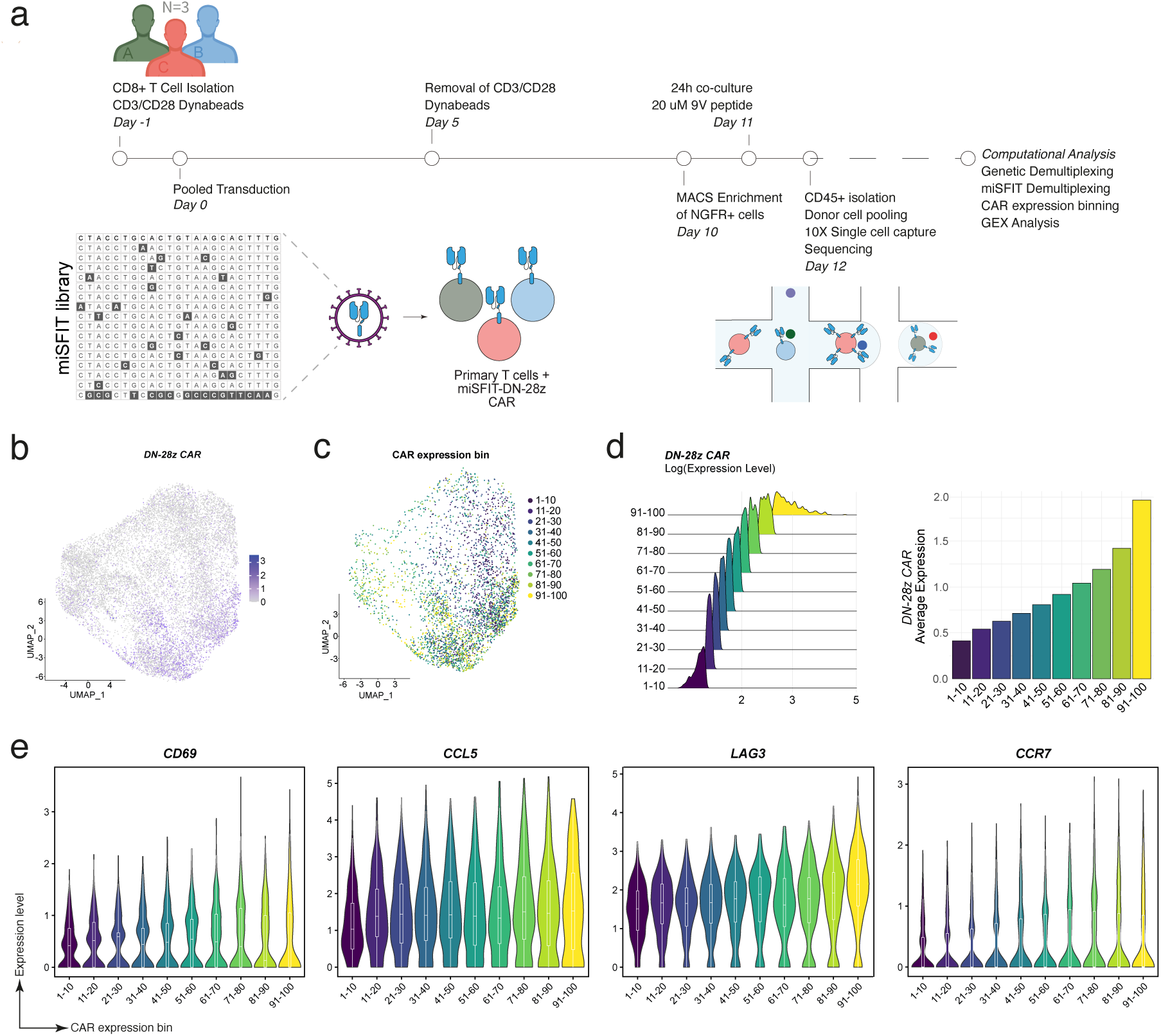
Dissecting CAR expression’s impact on the T cell transcriptome. **a)** Timeline and overview of scRNA-seq experiment, used to explore the influence of CAR levels on gene expression programmes. We created single-cell RNA-sequencing libraries from CD8+T cells transduced with a pool of lentivirus containing 19 DN-28z CAR miSFIT variants (*Methods and Materials*, **Extended Data** Figure 10-11). **b)** Uniform Manifold Approximation and Projection (UMAP) visualisation of cells captured in **(a)** (n=13,921 cells), coloured by *DN-28z CAR* expression level. **c)** UMAP of cells coloured by CAR expression decile bin. **d)** *DN-28z CAR* transcript expression distribution (left) and average expression for each CAR expression bin (right). **e)** Expression of representative marker genes for activation (*CD69*), effector (*CCL5*), exhaustion (*LAG3*), and memory (*CCR7*) plotted by CAR expression bin.

After scRNA-seq library preparation, sequencing, and pre-processing, we genetically demultiplexed cells originating from each of the three donors (**Extended Data** Figure 11). To annotate cells by the CAR-miSFIT variant they received, we performed targeted amplification and sequencing of miSFIT sequences along with their associated 10x barcodes (**Extended Data** Figure 12). After quality control filtering, we confidently assigned miSFIT sequences to 4,288 out of 13,921 single cells visualised on the UMAP plot (∼30% of cells, which aligns with previous tag recovery rates obtained by targeted tag amplification and sequencing) (**Extended Data** Figure 12d and **Figure 3b**).

We first sought to validate the tuning of the *DN-28z CAR* transgene. *DN-28z CAR* transcripts were detected in 27% of all cells (3,825 of 13,921 cells), with CAR expression levels emerging as a major driver of the overall transcriptional state, as evidenced by a clear gradient in CAR expression across the UMAP space **Figure 3b & c**). We quantified CAR expression for cells annotated by miSFIT variant and verified that miSFITs conferred stepwise control over expression levels (**Extended Data** Figure 12g. To quantify the statistical relationships between CAR expression and the abundance of various transcripts, we binned cells into CAR expression deciles (**Figure 3c & d**).

We examined the expression of marker genes for activation (*CD69*), effector function (*CCL5*), exhaustion (*LAG3*) and memory (*CCR7*) across the CAR expression bins. Each of these genes displayed a strong, positive, and continuous association with CAR expression level (**Figure 3e**), confirmed by robust and significant regression fitting between *DN-28z CAR* and T cell marker expression (**Extended Data** Figure 13). We did not detect significant derepression of endogenous miR-17-92 target genes across CAR expression bins (**Extended Data** Figure 14), verifying that the CAR miSFIT constructs do not act like miRNA sponges, consistent with our previous evaluation [14]. This confirms that the effects we observed on T cell marker genes were due to the tuning of the CAR transgene.

### 2.4 CAR expression correlates with a transcriptional signature of exhaustion

Next, we examined previously curated T cell activation, dysfunction, and memory modules [37] and calculated their enrichment score for each CAR expression bin (**Figure 4a** and **Extended Data** Figure 15). Dysfunction module scores rose with increasing CAR expression bins, revealing a continuous rather than switch-like association between CAR expression and dysfunction. In contrast, activation and memory scores both decreased slightly with increased CAR expression (**Figure 4a**). Taken together, these results provide insight into the relationships between T cell activation, dysfunction, and memory states and how they relate to CAR expression levels.

**Fig. 4:**
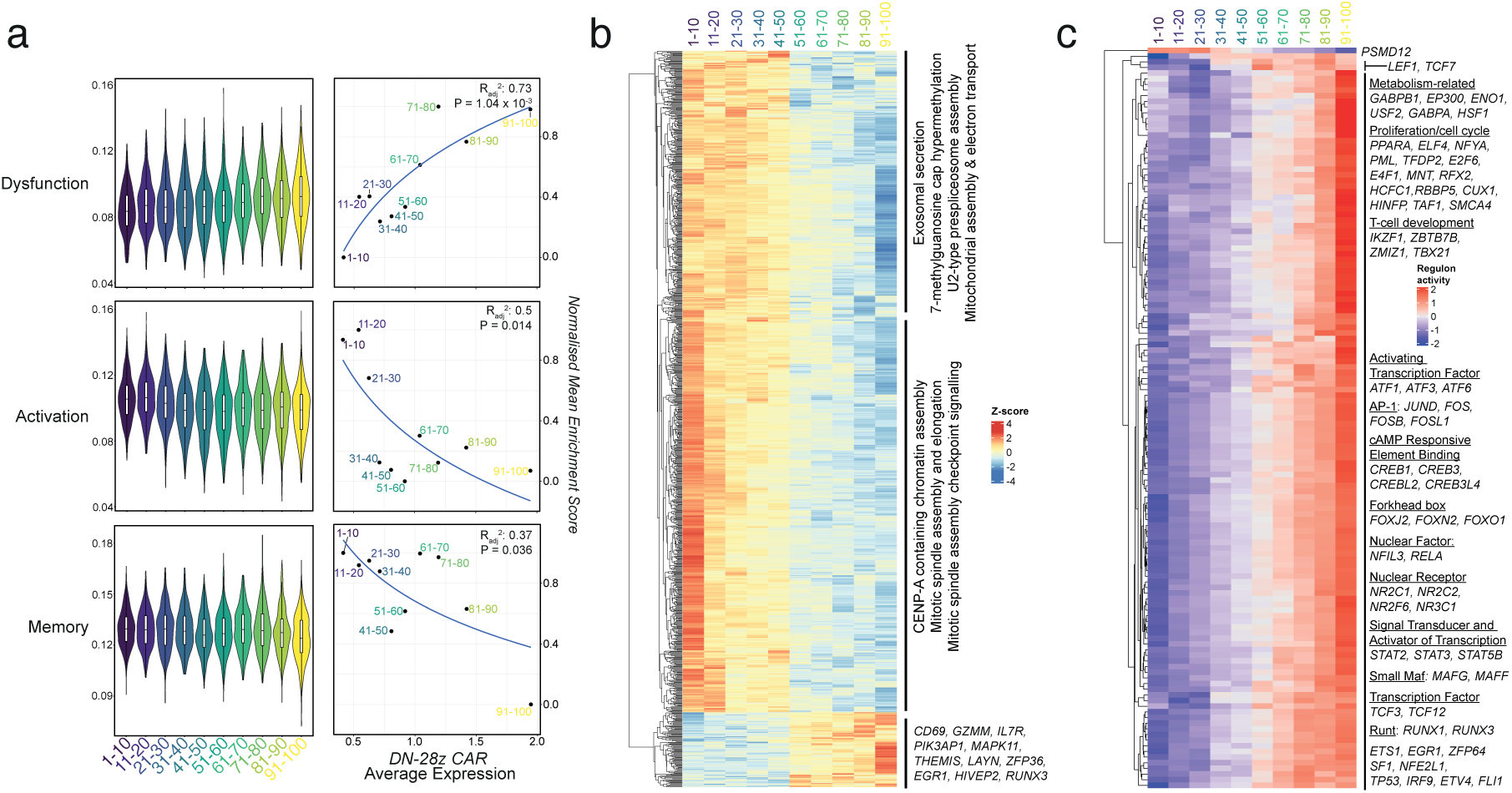
Supraphysiological CAR expression levels induces a global, dysfunctional transcriptional state. **a)** To investigate the relationship between CAR expression and T cell activation, dysfunction, and memory, we conducted gene set enrichment analysis (GSEA) using curated modules [37] on our CAR expression computational bins. Left: Violin plots activation, exhaustion, and memory modules per CAR expression bin. Right: Logarithmic regression fitting between normalised, average enrichment score and CAR expression bin. Enrichment scores were normalised against highest enrichment score for each module. R^2^ and P*_adj_* depict regression fit model parameters. **b)** Undertaking an unbiased exploration of CAR expression’s impact on the transcriptome, we calculated the mutual information (MI) [38, 39] between *DN-28z CAR* and the genes of our scRNA-seq atlas, filtering for genes with the highest mutual information. Heatmap and hierarchical clustering of genes identified with highest mutual information (0.693, n = 976 genes) between *DN-28z CAR* expression. Annotations are gene ontology (GO) themes and known T cell associated genes enriched in each expression module. **c)** SCENIC was used to identify transcription factor programmes impacted by CAR expression [73]. Heatmap and hierarchical clustering of regulons identified with SCENIC and subsequent filtering with highest mutual information between (n = 133 regulons) *DN-28z CAR* expression.

### 2.5 Revealing global, dysfunctional transcriptional programmes

After mapping the relationship between CAR expression and previously reported signatures of activation, memory, and exhaustion, we extended our analysis to a transcriptome-wide assessment. To identify genes that vary with CAR expression in a comprehensive and unbiased manner, we used three complementary approaches: mutual informational analysis, linear/logistic regression, and weighted correlation network analysis.

We first leveraged mutual information, a metric that quantifies statistical relatedness between variables [38, 39], to identify genes whose expression levels vary with CAR expression. We identified 976 genes that displayed high mutual information with CAR expression (MI = 0.693, **Figure 4b, Supplementary Table 1**), indicating the widespread impact that CAR expression levels have on the transcriptome. Hierarchical clustering of these genes revealed three modules with different expression pattern dynamics, including two modules of genes whose expression is negatively correlated (**Figure 4b**, upper clusters) and one where gene expression exhibits a positive correlation (**Figure 4b**, lower cluster) with CAR expression levels.

We used PantherDB [40, 41] to conduct gene ontology (GO) analysis and identify functions enriched in each module. The modules that were negatively correlated were enriched for numerous cellular processes, including gradually decreasing negative module was enriched for numerous cellular processes, including CENP-A containing chromatin assembly (Fold enrichment = 24.63, FDR = 1.26×10*^−^*^3^), mitotic spindle assembly checkpoint signalling (Fold enrichment = 13.13, FDR = 1.67×10*^−^*^4^), and exosomal secretion (Fold enrichment = 32.85, FDR = 3.17×10*^−^*^2^), **(Figure 4b, Supplementary Table 2 & 3)** [54–56]. These processes likely play a key role in clonal expansion during T cell activation and effector response. 7-methylguanosine cap hypermethylation (Fold enrichment = 27.38, FDR = 7.72×10*^−^*^4^), U2-type prespliceosome assembly (Fold enrichment = 18.25, FDR = 3.62×10*^−^*^7^) were additionally enriched, indicating an impairment in RNA-processing with high CAR expression. Finally, mitochondrial assembly (Fold enrichment = 13.42, FDR = 3.57×10*^−^*^12^) and electron transport (Fold enrichment = 15.24, FDR = 7.98×10*^−^*^11^) **(Figure 4b, Supplementary Table 2)** were identified, suggesting that metabolic processes and energy generation become impaired with increasing CAR expression levels. Indeed, expression of several transcripts related to oxidative phosphorylation, which include genes encoding all 5 complexes of the electron transport chain [47], exhibited a gradual reduction in expression with increasing CAR levels **(Extended Data** Figure 16a), underscoring their potential functional relevance in T cell dysfunction in high CAR-expressing T cells.

The module positively correlated with CAR levels was not enriched for any GO terms. However, this cluster included known factors involved in T cell activation, memory, and exhaustion, such as activation marker *CD69*, effector molecule *GZMM*, memory marker *IL7R* and T cell exhaustion marker *LAYN* [67], altogether underscoring the significance of CAR levels in producing widespread transcriptional changes that affect T cell phenotypes. Additionally, this cluster contained transcripts encoding for transcription factors involved in T cell development (e.g.,*THEMIS* [61–63] and *RUNX3* [69, 70]).

Next, we used linear and logistic regression analysis against the *DN-28z CAR* transcript and uniquely identified 155 and 89 additional genes identified, respectively (**Extended Data** Figure 16b**, Supplementary Table 4**). Gene ontology analysis of these newly identified genes revealed translation (Fold enrichment = 6.56, P*_adj_* = 9.10×10*^−^*^3^), mitochondrial translation (Fold enrichment = 9.51, P*_adj_* = 2.35×10*^−^*^2^), and nicotinamide nucleotide (e.g., precursor NAD^+^) metabolic process (Fold enrichment = 8.70, P*_adj_* = 4.46×10*^−^*^2^). These processes underscore CAR expression’s perturbation on foundational biological processes, widening its scope to protein expression and reiterating metabolic dysfunction.

Finally, we applied weighted correlation network analysis (WGCNA) to identify modules of highly correlated genes [107]; that varied similarly with changes in CAR expression. WGCNA identified 5,066 genes influenced by CAR expression levels (**Extended Data** Figure 16c **& d and Supplementary Table 5**). This expands the total number of genes affected by CAR expression to 6,043 genes (**Extended Data** Figure 16b), underscoring the global impact CAR expression has on the transcriptome. GO analysis of these newly identified genes revealed further biological processes (**Supplementary Tables 6-8**), including regulation of telomerase RNA localisation to Cajal body (Fold enrichment = 4.25, P*_adj_* = 4.29×10*^−^*^2^) and ribosomal small subunit assembly (Fold enrichment = 23.47, P*_adj_*= 2.74×10*^−^*^3^) (**Extended Data** Figure 16**, Supplementary Tables 6-8**). These findings demonstrate how high CAR expression levels perturb the biological processes essential for telomerase function [115] and protein synthesis.

Canonical glycolysis (Fold enrichment = 11.9, P*_adj_* = 5.06×10*^−^*^3^) and glycolytic process (Fold enrichment = 7.98, P*_adj_*= 4.93×10*^−^*^5^) were biological processes enriched for WGCNA module containing genes positively correlated with CAR expression. This suggests that increasing CAR expression results in a reliance on glycolysis for energy generation. Visualising component genes of canonical glycolysis confirms this, as genes encoding key glycolytic enzymes (*PGK1, HK1, HK2, ENO1, ENO2, ENO3, and TPI1*) showed increased expression levels with higher CAR expression bins (**Extended Data** Figure 16e). Glycolysis has been associated with terminally differentiated T cells and an impaired ability to form memory subtypes [73].

As transcription factors are core regulators of transcriptional programmes within cells, we used SCENIC package and mutual information (MI) to identify regulons (transcription factors and their downstream targets) influenced by CAR expression [74]. This revealed 132 regulons whose activity scaled with increasing CAR expression (**Figure 4c, Supplementary Table 9**). Regulons identified included those related to T-cell development, regulation, and activation, such as T-cell factor/lymphoid enhancer factor (TCF/LEF), signal transducer and activator of transcription (STAT), RUNX, cAMP responsive element binding (CREB), and AP-1 family members. Regulons with established roles in T cell dysfunction were also positively associated with increasing CAR expression, including FOXO1 [75] and TBX21 [76, 77] (**Figure 4c**). We also identified regulons with cellular metabolic functions, such as GABPA and GABPB1 (subunits of NRF-2) involved in mitochondrial biogenesis [78], PPARA controlling regulation of oxidative phosphorylation and fatty acid oxidation [79], and ENO1 driving glycolysis. These altered transcription factor programmes sit upstream of identified altered metabolic processes (**Figure 4b**, **Extended Data** Figure 16a **& e**). Regulons with reported roles in cell cycle regulation and proliferation were also positively associated with CAR expression, including E2F6 [80], PML [81], HCFC1 [82, 83]. Splicing factor 1, SF1, which has roles in spliceosome assembly and function [84], was also identified, complementing our earlier findings (**Figure 4b**).

The majority of the regulons identified by MI had a linear relationship with CAR expression (120/133 regulons had R^2^ ≥ 0.8 and P*_adj_ <* 0.05 for linear regression, **Extended Data** Figure 17). This shows that T cell transcriptional state changes in a continuous, rather than switch-like manner with increasing CAR expression.

Overall, these findings shed light on how multiple fundamental and essential biological processes become impaired with increasing CAR transgene levels, underscoring the global dysfunctional impact supraphysiological CAR expression has on the transcriptome. We detected a range of CAR expression (*<*30th percentile) where the gene expression programmes were only modestly impacted (**Figure 4b**, **Extended Data** Figure 16). This range is consistent with our findings of unperturbed gene modules (**Figure 4a**).

## 3 Discussion

Here, we have developed a system to fine-tune CAR transgene and used it to quantify the significance of CAR expression levels in modulating multiple T cell functions, including activation, cytokine production, and killing. Through single-cell RNA sequencing, we identified transcriptional dysregulation of fundamental biological processes through which high CAR expression impairs T cell phenotypes. Importantly, our findings demonstrate that reducing CAR expression can mitigate T cell dysfunction state and potentially enhance therapeutic efficacy.

Toxic side effects, such as cytokine release syndrome (CRS) and neurotoxicity (ICANS), affect a large number of patients undergoing CAR-T cell therapy, posing a barrier for widespread clinical application of these treatments [85, 118]. One of the factors contributing to such toxicities is the hyperactivation of CAR-T cells, which leads to excessive proliferation and cytokine-mediated systemic inflammatory responses [3, 4], further causing overreaction of the immune system [86]. On the other hand, the therapeutic efficacy of CAR-T cells depends on robust T cell activation and cytokine secretion. Thus, CAR-T cell activation must be carefully balanced within a therapeutic window to ensure efficacy while minimizing toxicity. Using miSFITs, we demonstrated that tuned CAR expression controlled both the magnitude of T cell activation and the proportion of activated cells (**Figure 1c & d**), consistent with previous findings [27]. miSFITs allows for refined control of CAR expression and T cell activation, providing an easy-to-implement approach for future studies addressing CAR-T toxicity.

An outstanding challenge of CAR-T therapy is on-target, off-tumour toxicity (OTOT). Indeed, in current FDA-approved therapies, B-cell aplasia is an expected side effect that requires clinical intervention [87]. Additionally, OTOT has significantly limited the successful application of CAR-T therapy to solid tumours. In large part, this issue stems from target antigen selection, where candidate tumour-associated antigens (e.g., EGFR and HER2) are expressed in both healthy and cancerous tissues [3, 5]. However, cancerous tissues generally express these antigens at much higher levels [88]. We observed that CAR expression levels are a key determinant of the target antigen dose required to initiate a T cell response (**Figure 1e**). This suggests that tuning CAR expression could be exploited to control the antigen density threshold required for robust T cell activation and cytotoxicity, offering a promising strategy to increase specificity in targeting cancer cells.

T cell dysfunction has severely limited CAR-T therapies and remains a challenge for achieving durable patient responses [2, 33, 89, 90]. We demonstrated how dysfunctional T cell phenotypes were enhanced in direct correlation with CAR expression, without improving activation or memory signatures (**Figure 4a**). Our unbiased analyses identified biological processes that became progressively dysregulated as CAR levels increased, including RNA processing, telomere maintenance, and metabolism. We also identified various processes associated with dysfunctional T cell activation and cytotoxicity, such as exosomal secretion. Exosomes have a role in targeted cell killing through ‘lethal hit delivery’ of granzymes and perforin [42], and exosomes originating from CAR T cells were recently shown to exhibit potent and specific anti-tumour efficacy *in vivo* [43]. Therefore, reduced exosomal secretion could hamper activity in T cells expressing supraphysiological levels of CAR. Similarly, dysregulation of 7-methylguanosine cap hypermethylation and U2-type prespliceosome [44–46] may reflect a dysfunction in RNA processing. In a similar vein, a perturbation in ribosome assembly and translation may reflect impaired protein synthesis. Together, impacting the global regulation of gene expression. Furthermore, gradual decrease in expression of genes involved in mitochondrial machinery and oxidative phosphorylation, as well as other cellular metabolic functions underscore the critical role of altered metabolism in causing T cell dysfunction, consistent with previous observations [48–53, 127]. Future studies are needed to dissect the mechanistic basis, dynamics and biological significance that dysregulation of these processes have on T cell phenotypes and CAR-T cell efficacy.

Our study utilised a second-generation CAR with a CD28 co-stimulatory domain. Given previous observations establishing CAR domains dictating CAR-T cell identity and function (e.g., phenotype, sensitivity, metabolic pathway preference, signalling kinetics and strength, etc.) [96, 113, 114], assessing the generalizability of our observations to other CAR designs would an insightful next step.

Low CAR expression (**Figure 4**) mitigated transcriptional T cell dysfunction whilst leaving activation and memory minimally impacted. This level correlated with the activity of regulons and core transcription factors associated with the biological processes altered during overactivation of CAR-T cells and T cell exhaustion (**Figure 4** and **Extended Data** Figure 16). Our findings provide further insight into T cell dysfunction.

FDA-approved CAR-T therapies are currently manufactured using gamma retroviral and lentiviral vectors. The miSFIT elements used in our study are fully compatible with these approved methods and are simpler to implement in the near term compared to targeted knock-in-based approaches for controlling CAR expression levels. Thus, integrating the miSFIT system with current CAR therapies provides a simple, straightforward, and clinically tractable method with the potential to achieve fine control over T cell state and function.

## 4 Methods

### 4.1 Generation of DN-28z miSFIT library

We generated DN-28z CAR miSFIT lentiviral expression vectors using standard restriction cloning methods. We used pPD-1-miSFIT-4x (Addgene plasmid #124678) as the parent vector. The DN-28z CAR was amplified from the pLEX-307-DN-SECOND-GEN-pDONR221 [94] using primers DN28zCAR-F and DN28zCAR-R. We digested this PCR product, as well as the destination vector with SbfI and NheI and ligated them using T4 DNA ligase (NEB). We transformed each ligation product into Subcloning Efficiency DH5a Competent Cells (Thermo Fisher) following manufacturer’s instructions. Transformants were plated overnight at 37*^◦^*C on 24.5cm^2^ ampicillin-treated LB agar plates. Each plasmid was recovered using the QIAprep Spin Miniprep kit (QIAGEN) and validated successful cloning using Sanger Sequencing with primers LentiBB-CMV-seq and LentiBB-SV40pA-seq, as seen in **8.12**.

For other miSFIT variants, we linearised and removed the 4x Perfect miSFIT from the validated plasmid using NheI and AgeI (NEB). We then generated other MRE inserts by synthesizing and annealing overlapping oligonucleotides comprising the MRE sequence and compatible sticky overhangs. We ligated the insert to the linearised vector for 1 hour at room temperature using T4 DNA Ligase (NEB) and transformed the ligation product into Subcloning Efficiency DH5a Competent Cells (Thermo Fisher) and proceeded with the previously described transformation and purification protocols. We validated successful cloning using Sanger Sequencing with primer LentiBB-SV40pA-seq.

To have a negative transduction control that could be detected with a miSFIT sequence, we cloned an empty expression vector, referred to as Empty-MRE. We used an online Random DNA Sequence generator to generate a DNA sequence of equal length to the DN-28z CAR gene (http://www.faculty.ucr.edu/~mmaduro/random.htm). We manually inspected this sequence, mutated transcription start sites and encoded frequent STOP codons throughout the open reading frame.We ordered it as gBlock Gene Fragment (IDT). We linearised and removed the NY-ESO-1 DN-28z CAR sequence from the NY-ESO-1-2ndGenCAR-WT-Mir17RE-1x plasmid by digesting with restriction enzymes SbfI (NEB) and NheI (NEB). We inserted the gBlock with the linearised plasmid using the In-Fusion HD Cloning System (Takara Clontech), following the manufacturer’s instructions. We followed the previously described transformation and plasmid purification protocols. We validated successful cloning using Sanger Sequencing with primers LentiBB-CMV-seq and LentiBB-SV40pA-seq. We then linearised and removed the 1x Perfect miSFIT sequence from this plasmid using restriction enzymes NheI and AgeI (NEB). A new scrambled sequence MRE insert was generated by synthesizing and annealing synthetic, overlapping oligonucleotides comprising the MRE and appropriate sticky overhangs. We followed the previously described ligation, transformation, and validation protocols for MRE insert introduction.

### 4.2 miRNA quantification

We used miRNeasy Mini kit (Qiagen) to extract miRNA from HEK293T and untransduced Jurkats. miRNA was then reverse transcribed using a target-specific structured RT primer for miR-17 and RNU6b, which serves as reference, housekeeping control, (TaqMan MicroRNA Assay ID 002308, 001093, Thermo Fisher) and Taqman MicroRNA Reverse Transcription Kit (Thermo Fisher) from 10ng total RNA input, following the manufacturers’ instructions. We performed RT-qPCR using target-specific RT primers (Thermo Fisher) and TaqMan Universal Master Mix II, no UNG (Thermo Fisher) on a CFX384 real-time system (Bio-Rad). The ΔΔCt method was used to compare expression miR-17 expression of Jurkat to HEK293T by comparing the Ct of miR-17 to that of RNU6b for each sample replicate.

### 4.3 Cell lines

HEK293T (CRL-3216) cells were obtained from American Type Culture Collection (ATCC) and 293T LentiX cells were purchased from Clontech. CHO-K1+hCD58+HLA-A2 and Jurkat-NF-*κ*B-eGFP were a gift from Omer Dushek (University of Oxford). HEK293T, LentiX, and CHO-K1 cells were cultured in DMEM high glucose (Gibco) supplemented with 10% FBS, 1 mM L-glutamine, 25 mM HEPES, 100 U ml-1 penicillin/streptomycin, and 1 mM pyruvate. Primary human T cells were cultured in CTL medium consisting of RPMI-1640 supplemented with 10% human serum, 2 mM L-glutamine, 25 mM HEPES, 100 U ml-1 penicillin/streptomycin and 50 µM *β*-mercaptoethanol. All cells were cultured at 37°C and 5% CO2 and frequently tested for the absence of mycoplasma.

HEK-293T cells were grown in Dulbecco’s modified Eagle’s medium (DMEM, Gibco) supplemented with 10% FBS (GIBCO), referred to as complete growth media. Cells were seeded in 24 well plates, 24 hours prior to transfection, allowing them to reach 85-90% confluency on transfection day. On the day of transfection, complete growth media was replaced with DMEM, 2% FBS, referred to as transfection media.

For DN-28z CAR transcript abundance tests and to create polyclonal cell lines, we used a previously developed Jurkat-NF-kB-eGFP reporter cell line [91]. Jurkats were grown in Roswell Park Memorial Institute 1640 with GlutaMAX medium (RPMI, Gibco) supplemented with 10% FBS (Gibco).

### 4.4 Lentiviral transduction of Jurkat reporter cell lines and primary T cells

To produce lentiviral particles in HEK-293T cells, we co-transfected each lentiviral transfer vector with pCMV-dR8.91 or psPAX2 and pMD2.G at a ratio of 1.5:1:1 using Polyethylenimine (PEI, 1 mg/mL, Sigma-Aldrich). After 24 hours, we exchanged the transfection media (DMEM (Gibco), 2% FBS (Gibco)) with full media (DMEM (Gibco), 10% FBS(Gibco)). We collected and filtered (0.22*µ*M filter, Millipore) viral supernatant 24 hours later and stored it at −80°C until transduction.

For DN-28z CAR RNA transcript measurement, an optimised pool of lentivirus encoding the entirety of DN-28z CAR miSFIT library was generated. Lentivirus for each DN-28z CAR miSFIT was separately generated as described above and respective functional viral titres were determined. Lentivirus was pooled for equal functional titers between each miSFIT variant. This was to ensure each CAR-miSFIT construct would transduce at similar rates, giving rise to a population of cells in which each miSFIT would be similarly represented (1x Perfect transduced into 500 cells, 2x Perfect transduced into 460 cells, . . ., V15 transduced into 480 cells, and Scramble transduced into 520 cells). Jurkat-NF-kB-eGFP cells were transduced with this lentiviral pool at a multiplicity of infection (MOI) less than 0.25 (below 10% transduced).

To generate miSFIT cell lines, Jurkat-NF-kB-eGFP cells were transduced at a low MOI (below 10% transduced), incubated for 5 days, and selected stably transduced cells by FACS, sorting on NGFR positive cells, using BD FACSAria Fusion.

### 4.5 gDNA and cDNA library preparation and high-throughput sequencing

We used the All Prep DNA/RNA Mini kit (Qiagen) to simultaneously extract genomic DNA (gDNA) and mRNA from Jurkats transduced with the DN-28z CAR-miSFIT library. After performing a genomic DNA wipe-out step (TURBO DNA free, Thermo Fisher), cDNA was generated from mRNA using the QuantiTect Reverse Transcription kit (Qiagen) following the manufacturer’s instructions. To create amplicon libraries for high-throughput sequencing, the miSFIT sequence and a short flanking region were PCR amplified using the primers bi-dir-Miseq-F and bi-dir-Miseq-R. For gDNA and cDNA, we used Phusion High-Fidelity PCR Master Mix with GC Buffer (NEB) and the following cycling conditions: initial denaturation (98°C for 3 minutes), 25 amplification cycles (98°C for 10s, 60°C for 12s, 72°C for 12s), and final extension (72°C for 10 minutes). PCR products were gel-purified using the QIAquick Gel Extraction Kit (Qiagen). The recovered product was diluted between 10 and 30 fold depending on band intensity. To make amplicon libraries compatible with Illumina machines, we performed a second PCR to append TruSeq index sequences and p5/p7 adapters to each amplicon. We used a dual barcoding strategy where a unique combination of forward and reverse index primers were assigned to each biological sample. We performed the PCRs with Phusion High-Fidelity PCR Master Mix with GC Buffer (NEB), using the following cycling conditions: initial denaturation (98°C for 30 seconds), 13 amplification cycles (98°C for 10s, 62°C for 10s, 72°C for 10s), and final extension (72°C for 5 minutes). We used Agencourt AMPure XP beads (1X, Beckman Coulter) at 1X concentration to purify the amplicon libraries. The libraries were quantified using the Qubit dsDNA HS Assay Kit (Thermo Fisher). Samples were pooled and sequenced (150bp PE) on the MiSeq v3 (Illumina).

High-throughput sequencing data were analysed using R (version 4.2) and all scripts are available via Github repository: https://github.com/ahsr-cell/Ramos2024. After inspecting the quality of sequencing data with FastQC (Babraham Institute), we used the Biostrings package (version 2.65.3) to trim reads down to the miSFIT sequence and subsequently count the occurrence of each type of miSFIT in all amplicon libraries. We calculated the transcript abundance for each variant by dividing its read count in the cDNA library to its read count in the respective gDNA library.

### 4.6 Flow cytometry and fluorescence activated cell sorting

All flow-cytometry experiments were performed on the BD LSR Fortessa Analyzer, BD FACSAria Fusion, or BD FACSAriaIII (BD Biosciences) and data were analysed using FlowJo (Version 10.8.1). For experiments requiring antibody staining, cells were washed with FACS buffer (PBS (Gibco) with 5% FBS (Gibco)) before and after staining. For sorting experiments, we used BD FACSAria Fusion or BD FACSAriaIII (BD Biosciences) with a 100*µ*M sorting chip. Untransduced cells and FMO controls were used to set sorting and analysis gates.

### 4.7 CD8+ T cell isolation, transduction, and expansion for T cell activation studies

CD8 T cell isolation and transduction was previously described and summarised below [93, 94]. Ethical approval was obtained by the Genetic Modification Safety Committee (IMM249) and Inter-Divisional Research Ethics Committee of the University of Oxford (R51997/RE001).

Human CD8+ T cells were isolated from four donor leukocyte cones purchased from the National Health Service’s (UK) Blood and Transplantation service. Isolation was performed using negative selection. Briefly, blood samples were incubated with Rosette-Sep Human CD8+ enrichment cocktail (Stemcell) at 150 *µ*l/ml for 20 minutes. This was followed by a 3.1 fold dilution with PBS before layering on Ficoll Paque Plus (GE) at a 0.8:1.0 ficoll to sample ratio. Ficoll-Sample preparation was centrifuged at 1200g for 20 minutes at room temperature. Buffy coats were collected, washed, and isolated cells counted. Cells were resuspended in complete primary T cell media (RPMI (Thermo Fisher) supplemented with 10% FBS (Gibco), 100 Units/ml penicillin, 100 *µ*g/ml streptomycin (Gibco)) with 50 U/ml of IL-2 (PeproTech) and CD3/CD28 Human T-activator Dynabeads (Thermo Fisher) at a 1:1 bead to cell ratio. Isolated human CD8+ T cells were cultured at 37°C and 5% CO2.

On the following day, 1 million cells were transduced using lentivirus titrated for a low multiplicity of infection (below 15% transduced). On days 2 and 4 post-transduction, 1 ml of media was exchanged, and IL-2 was added to a final concentration of 50 U/ml. Dynabeads were magnetically removed on day 5 post-transduction. CAR T cells were further cultured at a density of 1 million cells/ml and supplemented with 50 U/ml IL-2 every other day.

Day 8 post-transduction, a subpopulation of CAR Ts were harvested to measure DN-28z CAR expression levels. 500,000 CAR T cells were harvested, washed with FACS buffer (PBS (Gibco) + 5% FBS (Gibco)) before and after staining. T cells were stained with NY-ESO-1 9V PE–conjugated tetramer (produced in-house in the Dushek laboratory using refolded HLA*A0201 with NY-ESO-1 9V and streptavidin–PE [Bio-Rad AbD Serotec or BioLegend]). Cells were stained with antibody, diluted in FACS buffer, by incubating with cells for 30 minutes at 4°C. CAR expression was measured using flow cytometry. In downstream analysis, CAR expression was calculated by geometric median fluorescence intensity of NY-ESO-1 9V-PE. CAR expression was normalised on a donor basis by dividing by Scramble control expression.

### 4.8 Synthetic activation co-culture and measurement

To study the effects of tuned CAR expression on T-cell activation, we co-cultured DN-28z CAR-miSFIT T cells with target cells, CHO-K1+hCD58+HLA-A2.

Target cells used in these co-cultures were CHO-K1+hCD58+HLA-A2 cells. This is a variant of Chinese Hamster Ovary K1 (CHO-K1) cells that were transduced to expression human CD58 and HLA-A*02. HLA-A*02, human leukocyte antigen A*02, is an MHC-I (major histocompatibility complex I) molecule that binds peptide fragments and displays them on the cell surface for recognition by T cells. CD58 is a cell adhesion molecule that functions to strengthen the binding of a T cells to its target cell by binding to CD2 on the T cell.

For Jurkat-NFkB reporter tests, 30,000 CHO-K1 cells were seeded in 96 well plates, 24 hours prior to co-culture. On the day of co-culture, DMEM, 10% was replaced with Opti-MEM containing serial diluted concentrations, ranging from 0 nM to 20 nM, of NY-ESO-1_157-165_ 9V peptide. CHO-K1 cells were incubated with 9V peptide for 90 minutes for peptide pulsing. 9V peptide medium was aspirated and DN-28z CAR miSFIT Jurkat cell lines were added to each well at a 2 CAR-T : 1 CHO-K1 ratio. Co-cultures were incubated for 12 hours and harvested using 0.05% Trypsin with EDTA (Thermo Fisher Scientific). Cells were washed with FACS buffer (PBS (Gibco) with 5% FBS (Gibco)) before and after antibody staining for NGFR (BD Biosciences). Cells were stained with antibody, diluted in FACS buffer, by incubating with cells for 30 minutes at 4°C. Cells were analyzed with flow cytometry.

For CD8+ T cell tests, CD8 T cells were isolated, transduced, expanded as described above. On day 10 post-transduction, T cells were enriched for transduced cells using magnetic activated cell sorting (MACS) and CD271 (LNGFR, low-affinity, Nerve Growth Factor Receptor) MicroBead kit (MiltenyiBiotec) according to manufacturer’s instructions. Briefly, cells were resuspended in MACS buffer (PBS (Gibco) + 2% FBS (Gibco), degassed before use) and magnetically labelled by incubating with CD271 microbeads (Miltenyi Biotec) for 15 minutes at 4°C. Cells were washed after staining and resuspended in MACS buffer. Cells were applied to LS columns (Miltenyi Biotec) on QuadroMACS separator magnet (Miltenyi Biotec). The column was washed 3x with MACS buffer before being removed from magnet, and cells were plunged out. Enriched CAR-T Cells were resuspended in complete primary T cell media without IL-2 supplementation and rested overnight.

The same day of MACS enrichment, 180,000 CHO-K1 cells were seeded in 24 well-plates. On Day 11, post-transduction, co-cultures were conducted as described above. However, only two peptide concentrations of NY-ESO-1_157-165_ 9V peptide were used: 0nM and 20,480nM. On Day 12, post-transduction, cells were washed with FACS buffer (PBS (Gibco) with 5% FBS (Gibco)) before and after ZombieGreen (BioLegend) and antibody staining for CD45-BV421 and CD69-AF647 (BioLegend). Cells were stained with ZombieGreen, diluted in PBS (Gibco), at room temperature for 30 minutes, following manufacturer’s instructions. Antibodies for CD45-BV421 and CD69-AF647 subsequently were added, diluted in FACS buffer, and incubated with cells for 30 minutes at 4°C. ZombieGreen served as a LIVE/DEAD stain. CD45 served to discern CAR Ts from any residual CHO-K1 cell. Cells were analysed with flow cytometry. In downstream analysis, CD69 expression was calculated by geometric median fluorescence intensity of CD69-AF647. CD69 expression was normalised on a donor basis by dividing by activated (e.g., 20,480nM) Scramble control expression. CD69 %+ was normalised on a donor basis by subtracting unactivated (e.g., 0nM) untransduced control %+.

### 4.9 *In vitro* functional assays

Functional tests, CD8 T cell isolation and transduction was previously described [95, 96] and summarised below.

To assess tonic signalling, lentivirus for a selection of DN-28z CAR miSFIT was generated and used to transduce Jurkat triple reporter cell line (reporting NF-kB, NFAT, and AP-1 activity, [92]), as described above. Post-transduction, cells were harvested, stained with Tetramer-NY-ESO-1-APC (manufactured by Fred Hutchinson Immune Monitoring Core) and NGFR-PE (Biolegend), washed once, and analysed by flow cytometry.

*Acquisition of T cells from healthy donors for functional tests* Healthy individuals *>*18 years-old were enrolled in a Fred Hutchinson Cancer Center (FHCC) Institutional Review Board-approved study for peripheral blood collection (IRB protocol 344). Informed consent was obtained from all enrollees. Researchers were provided donor age, nondescript donor ID number, and were blinded to all other personally-identifiable information about study participants. Peripheral blood mononuclear cells (PBMC) were isolated by density gradient using Lymphocyte Separation Media (Corning). CD8+ T cells were isolated using the EasySep Human CD8+ T Cell Isolation Kit (StemCell Technologies) in accordance with manufacturer’s instructions.

To prepare CAR T cells, LentiX cells were transiently transfected with the CAR vector, psPAX2 and pMD2.G. One day later (day 1), primary T cells were activated using Dynabeads Human T-Activator CD3/CD28 (ThermoFisher) at a 3:1 bead to T cell ratio and cultured in CTL supplemented with 50 U ml-1 IL-2 (Prometheus). The next day (day 2), lentiviral supernatant was harvested from LentiX cells, filtered using a 0.45 µm PES filter, and added to activated T cells. Polybrene was added to reach a final concentration of 4.4 mg ml-1, and cells were spinoculated at 800g for 90 minutes at 32°C. Viral supernatant was replaced 8 hours later with fresh CTL medium supplemented with 50 IU 8 ml-1 IL-2. Half-media changes were then performed every 48 hours using CTL supplemented with 50 IU ml-1 IL-2. Dynabeads were removed on day 6, CD8+EGFRt+ transduced T cells were FACS-sorted on day 9, and purified CD8+EGFRt+ cells were rapidly expanded using 30 ng ml-1 purified OKT3 (30 ng/ml), *γ*-irradiated allogenic lymphoblastoid cell line (LCL, 8000 rad), and *γ*-irradiated (3500 rad) allogeneic PBMC at a LCL to T cell ratio of 100:1 and a PBMC to T cell ratio of 600:1. 50 IU ml-1 IL-2 was added on day 1, OKT3 was washed out on day 4, cultures were fed with fresh CTL medium supplemented with 50 IU ml-1 IL-2 every 2-3 days and resting T cells were assayed 11-12 days after stimulation. Non-transduced T cells (CD8+EGFRt-T cells that were not transduced with lentivirus) were cultured identically and used as negative controls for CAR T assays.

CAR T cells were co-cultured with T2 cells loaded with 10*µ*M NY-ESO-1 9V peptide. To measure cytokine secretion, cells were co-cultured at an 1:1 effector to target (E:T) cell ratio. Cytokine concentrations in cellular supernatant after 24 hours were quantified by human IFN-*γ* enzyme-linked immunosorbent assay (Thermo Fisher Scientific). For T cell killing, T2 cells were additionally incubated with ^51^Cr (PerkinElmer) for 16 hours at 37°C, washed with RPMI, and plated with 1,000 ^51^Cr-labeled target cells per well. CAR-T cells were added in triplicate at E:T ratios of 30/10/3/1:1 and incubated at 37°C, 5% CO_2_ for four hours. Supernatants were then harvested for *γ*-counting, and specific lysis was calculated by comparing counts to standardized wells in which target cells were lysed with a NP40-based soap solution. T cell proliferation was quantified by staining CAR T cells with 1*µ*M solution of CellTrace Violet dye (Thermo Fisher scientific) prior to co-culture with T2 cells at a 1:2 E:T ratio. After 72 hours, cells were harvested, stained with fluorescently labelled anti-human CD8 antibody, washed once, and analysed by flow cytometry. The frequency of divided cells was calculated by drawing a “% Undivided” gate on the undivided peak in negative control samples and then setting a “% Divided” that bordered the first “% Undivided” gate. Activation induced cell death (AICD) was quantified via staining with SYTOX Green (Thermo Fisher Scientific) and Caspase-3/7 (Thermo Fisher Scientific) after a 24h co-culture at a 1:2 E:T ratio. To assess sensitivity, we measured phosphorylation of extracellular-signal-regulated kinases (pERK) and IFN-*γ* production by intracellular cytokine staining (ICS). To quantify pERK, well plates were coated in serial dilutions of avidin and NY-ESO-1 monomer-biotin before T cells were added. T cells incubated for 10 minutes before measuring pERK with flow cytometry. For ICS, T2 cells were loaded with serial dilutions of NY-ESO-1 9V peptide. After a 5h co-culture, IFN-*γ* was measured with flow cytometry.

### 4.10 Analysis of CAR expression, T cell activation, and function experiments

Prism version 10 (GraphPad Software) was used to plot data and calculate statistics. Precise statistical tests used are indicated in figure legends.

### 4.11 Single cell RNA sequencing

*RNA library preparation* To create a single cell expression atlas of fine-tuned CAR expression, an optimised pool of lentivirus encoding the entirety of DN-28z CAR miSFIT library was generated. As done previously, lentivirus for each DN-28z CAR miSFIT was individually generated and respective functional viral titer were determined. Lentivirus was pooled for equal functional titers between each miSFIT variant. We validated our pooling by conducting a test transduction on Jurkat T cells and validated the number of genomic integrations by deep sequencing (**Methods and Materials, Extended Data** Figure 10b). Using this optimised lentiviral pool, we transduced each donor’s T cells at a dose for an MOI of less than 0.25 (below 10% transduced) (**Extended Data** Figure 10c). As in previous transductions, this was done to avoid confounding effects on CAR expression level.

CD8+ T cells were isolated from three donors, as previously described above. On day 0, isolated T cells were transduced with an optimised pool of lentivirus generated as described above. Post-transduction, T cells were expanded. Day 10 post-transduction, T cells were MACS enriched, using CD271 MACS microbead kit (Miltenyi Biotec). To have untransduced, negative control T cells, MACS flow-through (containing unlabeled, untransduced cells) was spun down, counted, and spiked in to enriched T cell populations at 1:21 ratio. T cells were rested overnight.

The same day of MACS enrichment, 900,000 CHO-K1 cells were seeded in 6 well plates. Co-cultures were conducted as described above, being initiated Day 11 post-transduction. However, only 20,480nM peptide concentration of NY-ESO-1_157-165_ 9V peptide were used.

On Day 12 post-transduction, cells were washed with FACS buffer (PBS (Gibco) with 5% FBS (Gibco)) before and after ZombieGreen (BioLegend) and antibody staining for CD45-BV421 (BioLegend). Cells were stained with ZombieGreen, diluted in PBS (Gibco), at room temperature for 30 minutes, following manufacturer’s instructions. Cells were stained with CD45-BV421 antibody, diluted in FACS buffer, at 4°C for 30 minutes. CD45+ cells were selected by FACS using BD FACSAria III and BD FACSAria Fusion. CD45 served to discern CAR Ts from any residual CHO-K1 cell. Furthermore, selection on CD45 included both untransduced cells and enriched, CAR T cells. Post-FACS sorting, a subpopulation of each donor’s, FACS-sorted T cells were saved for downstream bulk RNA-sequencing library preparation.

Each donor’s FACS-sorted T cells was pooled at a 1:1:1 ratio. This pool was diluted and approximately 84,000 cells were superloaded across two Chromium 10X Chip channels (10X Genomics). 10X Chromium Next Single Cell 3’ v3.1 kit (10X Genomics) was used for single cell capture, cDNA synthesis, and single-cell RNA-sequencing (scRNA-seq) library construction, following manufacturer’s instructions. scRNA-seq library was quality-controlled using Agilent TapeStation and quantified using Qubit. scRNA-seq library was sequenced on NovaSeq 6000 S4 (150bp PE).

cDNA generated from the 10X single-cell reaction was used to create PCR amplicon libraries that captured the 10X cell barcode and miSFIT sequence. Primers 10X-CARmiSFIT-F and 10X-CARmiSFIT-R were used to PCR amplify the miSFIT sequence and 3’ captured 10X cDNA. We used NEBNext Ultra II Q5 Master Mix (NEB) and the following cycling conditions: initial denaturation (98°C for 30 seconds), 30 amplification cycles (98°C for 10s, 65°C for 12s, 72°C for 12s), and final extension (72°C for 10 minutes). Primers 10x-EmptymiSFIT-F and 10x-EmptymiSFIT-R were used to separately capture the Empty miSFIT. NEBNext Ultra II Q5 Master Mix (NEB) and the following cycling conditions: denaturation (98°C for 30 seconds), 25 amplification cycles (98°C for 10s, 69°C for 12s, 72°C for 12s), and final extension (72°C for 10 minutes). Gel electrophoresis was used to validate the size of the resulting PCR products.

These initial PCR products were gel-purified using the QIAquick Gel Extraction Kit (Qiagen). We further purified these products using Agencourt AMPure XP beads (1X, Beckman Coulter) at 1X concentration. To make amplicon libraries compatible with Illumina machines, We adopted the dual-barcoding strategy as previously described above.

To append on TruSeq index sequences and p5/p7 adapters to the 10X CAR-miSFIT libraries, We performed PCRs using TruSeq-501-10XCAR and TruSeq-701-10XCAR with NEBNext Ultra II Q5 Master Mix (NEB) and the following cycling conditions: denaturation (98°C for 30 seconds), 13 amplification cycles (98°C for 10s, 61°C for 12s, 72°C for 12s), and final extension (72°C for 10 minutes). For the Empty library, We used TruSeq-501 and TruSeq-701 with NEBNext Ultra II Q5 Master Mix (NEB) and the following cycling conditions: denaturation (98°C for 30 seconds), 13 amplification cycles (98°C for 10s, 61°C for 12s, 72°C for 12s), and final extension (72°C for 10 minutes).

We used Agencourt AMPure XP beads (1X, Beckman Coulter) at 1X concentration to purify the amplicon libraries. The libraries were quality-controlled using 2100 Bioanalzyer (Agilent) and quantified using the Qubit dsDNA HS Assay Kit (Thermo Fisher). The 10X-CAR-miSFIT library was pooled with the 10X-Empty-miSFIT library at a 1:21 ratio. The final library was sequenced (300bp PE) on the MiSeq v3 (Illumina).

For each T cell donor, separate bulk RNA-sequencing libraries were generated for downstream genetic demultiplexing. We used All Prep DNA/RNA Mini kit (Qiagen) to simultaneously extract genomic DNA (gDNA) and mRNA from 1×10^6^ FACS-sorted T cells. After a genomic DNA wipe-out step (TURBO DNA free, Thermo Fisher), we created RNA-sequencing libraries using SMART-Seq v4 Ultra Low Input RNA Kit (Takara Bio) and Nextera XT DNA Library (Illumina) following manufacturer’s guidelines. Libraries were QC using 2100 Bioanalzyer (Agilent) and quantified using Qubit (Thermo Fisher). Libraries were sequenced on NextSeq 500 (75bp PE).

#### Single cell RNA-seq computational analysis

For all RNA-sequencing analysis, we used a single custom human genome reference that featured the DN-28z CAR miSFIT transcript. We used Genome Reference Consortium Human Build 38 patch release 13 genome assembly (GRCh38.p13) and appended on the DN-28z CAR-miSFIT transcript using Cellranger’s ’mkref’ function. We refer to this custom genome as GRCh38.p13.DN28zmiSFIT. Single-cell RNA-seq fastq files were demultiplexed using Cellranger (v6.1.2 10X Genomics) and mapped to the previously described GRCh38.p13.DN28zmiSFIT genome. 20,171 estimated cells were obtained with a mean reads per cell of 141,178, median genes per cell of 2,653, median UMI counts per cell of 9,768, and total number of genes detected was 16,524. 96.2% of reads had valid barcodes with a Q30 of 94.7% and 94.1% of the reads mapped confidently to the human genome. Downstream analysis and visualization were carried out in R (v4.2) using Seurat (v4.1.1, [97]). Matrices were filtered to remove barcodes with fewer than 200 genes (nFeature RNA), more than 6000 genes expressed and carried a high percentage of UMIs from mitochondrial features (percent.mt, greater than 10%) (Figure 2.2a).

EmptyDrops (v1.17.1, [98]) was used on Seurat-filtered barcodes to identify and filter out empty droplets. scDblFinder (v1.11.3, [99]) was used on EmptyDrops-filtered barcodes to identify and filter out any doublets. Outstanding barcodes were then subjected to genetic demultiplexing and miSFIT demultiplexing pipelines.

Genetic demultiplexing was used to identify cells’ donor origin, as previously described [100]. In brief, bulk RNA sequencing libraries from each donor’s cells were separately created and sequenced. Donor bulk RNA-sequencing fastq files were quality-control checked with FastQC. Adaptor sequences were trimmed using TrimGalore (v0.6.5, Babraham Institute) and aligned reads to custom reference genome (GRCh38.p13.DN28zmiSFIT) using STAR (v2.7.3, [101]). Minor modifications were made to the variant calling pipeline to incorporate metadata into the BAM alignment files and produce a block gVCF file. GATK variant calling pipeline (v4.1.7) was applied to BAM alignment files to identify sequence variants. cellSNP (v0.3.1, [102]) and Vireo (v0.4, [103]) were used to genotype and demultiplex filtered barcodes.

10X-PCR amplicon library capturing 10X cell barcode, DN-28z CAR, and miSFIT sequences were analysed using R (version 4.2) and all scripts are available via Github repository: https://github.com/ahsr-cell/Ramos2024. After inspecting the quality of sequencing data with FastQC, We used ShortRead (v1.55, [104]), Biostrings package (v2.65.3), stringr (v1.4.1), and dplyr (v1.0.10) to write an algorithm of the following:

1. Searched for the Seurat QC-filtered 10X cell barcodes in Read 1 of the resulting PCR amplicon fastq files
2. Counted the number of times any miSFIT sequence appeared in the respective Read2
3. For a cell barcode with any detected miSFIT counts, considers the two miSFIT with the highest counts (referred to as M1, miSFIT with the highest, and M2, second highest, from hereon) and assigns a miSFIT based on the following requirements, designed to remove ambiguous cases and intra-patient multiplets:

(a) M1 does not equal M2
(b) M1 exceeds a minimum read count of 25 (**Extended Data** Figure 12b)
(c) The ratio of M1 to M2 exceeds minimum factor threshold of 3 (**Extended Data** Figure 12c)
(d) Should any of the above requirements be failed, the cell was unlabelled and the cell-barcode discarded

Donor identified and miSFIT identified barcodes were annotated into our single cell expression matrix. Cells that were not donor identified were filtered out, resulting in 13,921 barcodes used for analysis. Expression values for the total UMI counts per cell were log-normalized and scaled. Highly variable genes were computed before performing principal component analysis (using 50 principal components). Linear dimensional reduction was conducted (resolution = 0.5, dims = 1:50). For down-stream analysis, the single cell expression matrix was subsetted to only include cells with detected DN-28z CAR expression (**Figure 3b & c**), resulting in 3,825 cells being in the single cell expression matrix. Cells were subsequently ranked according to their percentile rank in *DN-28z CAR* expression. 10 bins were created based on DN-28z CAR percentile rank: 1-10, 11-20, 21-30, 31-40, 41-50, 51-60, 61-70, 71-80, 81-90, 91-100, and cells were binned and annotated within the single cells expression matrix accordingly (**Figure 3c & d**). These bins are hereafter referred to as CAR expression bins.

Gene expression levels were averaged according to CAR expression bin and used for subsequent analysis, referred to as averaged expression matrix. Cells were scored using the escape package (v1.7.0) using UCell algorithm [105] for dysfunction, effector, and na ϊ ve/memory gene signatures published by Singer and colleagues [37] Enrichment scores were averaged according to CAR expression bin. Enrichment scores were averaged according to CAR expression bin. Mathematical modelling was conducted using the stats package (v3.6.2) and drc package (v3.0-1) and visualized using the ggplot2 package (v3.3.6).

Regulons were computed with the SCENIC (Single-Cell rEgulatory Network Inference and Clustering) pipeline (v1.3.1 [73]) using CAR expressing cells. GENIE3 (v.1.19.0) was used to determine coexpression modules. hg38 transcription factor motifs provided by the Aerts lab were used to identify direct transcription factor motifs using Rcistarget (v1.18.2). Regulon activity was quantified in single cells using the Area Under Curve metric built into the pipeline, calculated by AUCell (v.1.19.1). Linear and logarithmic regression was conducted using the stats package (v4.3.0) using the averaged expression matrix. Genes were regressed against the DN-28z CAR transcript. Any genes with a regression model R^2^ above 0.9 were used for subsequent analysis. We used the infotheo package (v1.2.0.1) using the average expression matrix to calculate the mutual information of all genes against the DN-28z CAR. Genes with an MI value of 0.693 were used for subsequent analysis. Weighted correlation network analysis was conducted using WGCNA (v1.72-1, [107]) using the averaged expression matrix. A power of 8 was used for the adjacency function and a merge cut height of 0.25 (for high resolution) and 0.45 (for low resolution) was used for module adjustment and collapsing. We focused on the turquoise and brown modules from the high resolution analysis and the magenta module from the low resolution analysis. We conducted gene ontology analysis on the turquoise and brown modules of the high-resolution WGCNA analysis and the magenta module of the low-resolution analysis (**Extended Data** Figure 16c **& d**). PantherDB (release 17.0, [106]) was used to conduct Gene Ontology analysis. Overrepresentation analysis was conducted using Fisher’s Exact test, calculating for false discovery rate (FDR) or Binomial test with the Bonferroni correction for multiple testing. For this analysis, a minimum overrepresentation fold enrichment of 10 (for Fisher’s exact testing) or 4 (for Binomial testing) and cut-off of FDR or P*_adj_*-value less than 0.05 was used. Bubble plot visualisation was created using Python v3.11.8, pandas v2.2.0, seaborn v0.13.2, matplotlib v3.8.0, plotly v5.11.0, and numpy v1.26.2. All heatmaps were visualised using pheatmap v1.0.12.

### 4.12 Data availability

Raw and processed data generated in this study were submitted to GEO with the following accession number **GSE268299**. Source data for **Figures 1-2**, **Extended Data** Figures 1-10 are provided with the paper.

### 4.13 Oligonucleotides, peptides, and plasmids used in this study

Included in the following table are the oligonucleotides used in this study. All oligonucleotides were synthesized by Integrated DNA Technologies. Peptides were synthesised at a purity of *>*95% (Peptide Protein Research, UK). 9V refers to a peptide derived from NY-ESO_157–165_ (SLLMWITQV). Relevant plasmids described in this study are available from Addgene (https://www.addgene.org/TatjanaSauka-Spengler/).

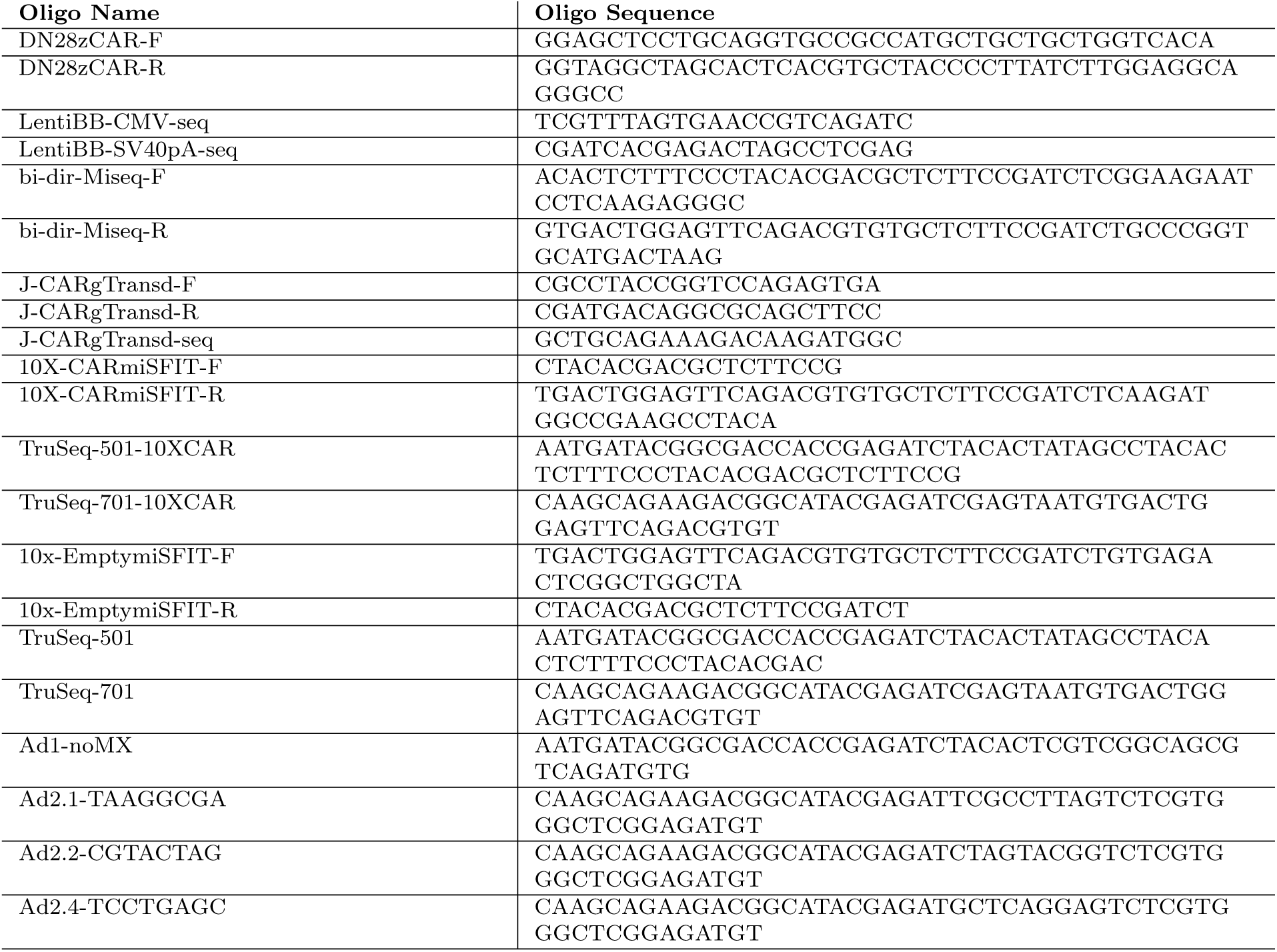

## Acknowledgements

We thank T. Baeumler for training. We acknowledge L. Overend for cloning of constructs. We thank the MRC WIMM Flow Cytometry, Single Cell, and Sequencing Facilities for providing training and technical support. We acknowledge Z. Hu and A. Rodriguez Delherbe for computational and technical advice. We thank Javier Carmona for his comments on the manuscript.

## 6 Funding

This work was supported by the Wellcome Trust Grant (ref 215615/Z/19/Z) and the institutional support from Stowers Institute for Medical Research to T.S.S.. A.S.R. was supported by the Sir Christopher Cox Junior Research Fellowship, provided by New College, University of Oxford. Y.S.M. is supported by the CancerCare Manitoba Foundation. S.S. is a Special Fellow of the Leukemia & Lymphoma Society (LLS career Development Program, Grant ID; 3405-21). O.D. is funded by the Wellcome Trust Senior Fellowships in Basic Biomedical Sciences (207537/Z/17/Z) and J.F.-S. is partly funded by the MRC project grant (MR/W031353/1)and Guy Newton Translational Grant (GN 05 (13)).

## 7 Author Contributions

A.S.R., T.S.S., T.A.F., and Y.S.M. conceptualised and designed the study. A.S.R. performed the majority of experiments and analysed and interpreted data. J.F.-S. and O.D. provided access and expertise in primary T cells and conducted transductions. E.Z.d.C. and O.P. participated in construct design. S.S. and A.R. generated and analysed functional data under the supervision of S.Riddell. S.Revale performed genetic demultiplexing data processing. T.S.S. and Y.S.M. supervised the project. A.S.R. wrote the original draft. A.S.R., Y.S.M., and T.S.S. reviewed and edited the draft. All other authors approved and provided feedback on the manuscript.

## 8 Conflicts of interest

S.Riddell is a cofounder and adviser to Lyell Immunopharma; has research funding from and intellectual property licensed to Lyell Immunopharma; was a cofounder of Juno Therapeutics; is an inventor of patents licensed to Juno Therapeutics; and served as an adviser to Juno Therapeutics and Adaptive Biotechnologies. Y.S.M. is an inventor on US patent 12018256.

## References

[1] June, C.H., Sadelain, M., 2018. Chimeric Antigen Receptor Therapy. New England Journal of Medicine 379, 64–73.

[2] Labanieh, L., Mackall, C.L., 2023. CAR immune cells: design principles, resistance and the next generation. Nature 614, 635–648.

[3] Brandt, L.J.B., Barnkob, M.B., Michaels, Y.S., Heiselberg, J., Barington, T., 2020. Emerging Approaches for Regulation and Control of CAR T Cells: A Mini Review. Frontiers in Immunology 11.

[4] Sterner, R.C., Sterner, R.M., 2021. CAR-T cell therapy: current limitations and potential strategies. Blood Cancer Journal 11.

[5] Flugel, C.L., Majzner, R.G., Krenciute, G., Dotti, G., Riddell, S.R., Wagner, D.L., Abou-El-Enein, M., 2023. Overcoming on-target, off-tumour toxicity of CAR T cell therapy for solid tumours. Nature Reviews Clinical Oncology 20, 49–62.

[6] Frigault, M.J., Lee, J., Basil, M.C., Carpenito, C., Motohashi, S., Scholler, J., Kawalekar, O.U., Guedan, S., Mcgettigan, S.E., Posey, A.D., Ang, S., Cooper, L.J.N., Platt, J.M., Johnson, F.B., Paulos, C.M., Zhao, Y., Kalos, M., Milone, M.C., June, C.H., 2015. Identification of Chimeric Antigen Receptors That Mediate Constitutive or Inducible Proliferation of T Cells. Cancer Immunology Research 3, 356–367.

[7] Gomes-Silva, D., Mukherjee, M., Srinivasan, M., Krenciute, G., Dakhova, O., Zheng, Y., Cabral, J.M.S., Rooney, C.M., Orange, J.S., Brenner, M.K., Mamonkin, M., 2017. Tonic 4-1BB Costimulation in Chimeric Antigen Receptors Impedes T Cell Survival and Is Vector-Dependent. Cell Reports 21, 17–26.

[8] Eyquem, J., Mansilla-Soto, J., Giavridis, T., Van Der Stegen, S.J.C., Hamieh, M., Cunanan, K.M., Odak, A., Gönen, M., Sadelain, M., 2017. Targeting a CAR to the TRAC locus with CRISPR/Cas9 enhances tumour rejection. Nature 543, 113–117.

[9] Ho JY, Wang L, Liu Y, Ba M, Yang J, Zhang X, Chen D, Lu P, Li J., 2021. Promoter usage regulating the surface density of CAR molecules may modulate the kinetics of CAR-T cells in vivo. Mol Ther Methods Clin Dev. 21, 237–246.

[10] Rodriguez-Marquez, P., Calleja-Cervantes, M. E., Serrano, G., Oliver-Caldes, A., Palacios-Berraquero, M. L., Martin-Mallo, A., Calviño, C., Español-Rego, M., Ceballos, C., Lozano, T., San Martin-Uriz, P., Vilas-Zornoza, A., Rodriguez-Diaz, S., Martinez-Turrillas, R., Jauregui, P., Alignani, D., Viguria, M. C., Redondo, M., Pascal, M., Martin-Antonio, B., . . . Prosper, F. 2022. CAR density influences antitumoral efficacy of BCMA CAR T cells and correlates with clinical outcome. Science advances, 8(39), eabo0514.

[11] Tristán-Manzano, M., Maldonado-Pérez, N., Justicia-Lirio, P., Muñoz, P., Cortijo-Gutiérrez, M., Pavlovic, K., Jiménez-Moreno, R., Nogueras, S., Carmona, M. D., Sánchez-Hernández, S., Aguilar-González, A., Castella, M., Juan, M., Marañón, M., Marchal, J. A., Benabdellah, K., Herrera, C., & Martin, F., 2022. Physiological lentiviral vectors for the generation of improved CAR-T cells. Molecular therapy oncolytics, 25, 335–349.

[12] Long, A.H., Haso, W.M., Shern, J.F., Wanhainen, K.M., Murgai, M., Ingaramo, M., Smith, J.P., Walker, A.J., Kohler, M.E., Venkateshwara, V.R., Kaplan, R.N., Patterson, G.H., Fry, T.J., Orentas, R.J., Mackall, C.L., 2015. 4-1BB costimulation ameliorates T cell exhaustion induced by tonic signaling of chimeric antigen receptors. Nature Medicine 21, 581–590.

[13] Calderon, H., Mamonkin, M., Guedan, S., 2020. Analysis of CAR-Mediated Tonic Signaling, in: Methods in Molecular Biology. Methods in Molecular Biology, pp. 223–236.

[14] Michaels, Y.S., Barnkob, M.B., Barbosa, H., Baeumler, T.A., Thompson, M.K., Andre, V., Colin-York, H., Fritzsche, M., Gileadi, U., Sheppard, H.M., Knapp, D.J.H.F., Milne, T.A., Cerundolo, V., Fulga, T.A., 2019. Precise tuning of gene expression levels in mammalian cells. Nature Communications 10.

[15] Prochazka, L., Michaels, Y.S., Lau, C., Jones, R.D., Siu, M., Yin, T., Wu, D., Jang, E., Vázquez-Cantú, M., Gilbert, P.M., Kaul, H., Benenson, Y., Zandstra, P.W., 2022. Synthetic gene circuits for cell state detection and protein tuning in human pluripotent stem cells. Molecular Systems Biology 18.

[16] Liu, W., Saito, Y., Jackson, J., Bhowmick, R., Kanemaki, M. T., Vindigni, A., & Cortez, D. 2023. RAD51 bypasses the CMG helicase to promote replication fork reversal. Science, 380(6643), 382–387.

[17] Landgraf, P., Rusu, M., Sheridan, R., Sewer, A., Iovino, N., Aravin, A., Pfeffer, S., Rice, A., Kamphorst, A.O., Landthaler, M., Lin, C., Socci, N.D., Hermida, L., Fulci, V., Chiaretti, S., Foà, R., Schliwka, J., Fuchs, U., Novosel, A., Müller, R.-U., Schermer, B., Bissels, U., Inman, J., Phan, Q., Chien, M., Weir, D.B., Choksi, R., De Vita, G., Frezzetti, D., Trompeter, H.-I., Hornung, V., Teng, G., Hartmann, G., Palkovits, M., Di Lauro, R., Wernet, P., Macino, G., Rogler, C.E., Nagle, J.W., Ju, J., Papavasiliou, F.N., Benzing, T., Lichter, P., Tam, W., Brownstein, M.J., Bosio, A., Borkhardt, A., Russo, J.J., Sander, C., Zavolan, M., Tuschl, T., 2007. A Mammalian microRNA Expression Atlas Based on Small RNA Library Sequencing. Cell 129, 1401–1414.

[18] Panwar, B., Omenn, G.S., Guan, Y., 2017. miRmine: a database of human miRNA expression profiles. Bioinformatics 33, 1554–1560.

[19] Thomas, R., Al-Khadairi, G., Roelands, J., Hendrickx, W., Dermime, S., Bedognetti, D., Decock, J., 2018. NY-ESO-1 Based Immunotherapy of Cancer: Current Perspectives. Frontiers in Immunology 9.

[20] Maus, M.V., Plotkin, J., Jakka, G., Stewart-Jones, G., Rivière, I., Merghoub, T., Wolchok, J., Renner, C., Sadelain, M., 2016. An MHC-restricted antibody-based chimeric antigen receptor requires TCR-like affinity to maintain antigen specificity. Molecular Therapy - Oncolytics 3, 16023.

[21] Abraham, R.T., Weiss, A., 2004. Jurkat T cells and development of the T-cell receptor signalling paradigm. Nature Reviews Immunology 4, 301–308.

[22] Smith-Garvin, J.E., Koretzky, G.A., Jordan, M.S., 2009. T Cell Activation. Annual Review of Immunology 27, 591–619.

[23] Vallabhapurapu, S., Karin, M., 2009. Regulation and Function of NF-*κ*B Transcription Factors in the Immune System. Annual Review of Immunology 27, 693–733.

[24] Xu, X., Nagarajan, H., Lewis, N.E., Pan, S., Cai, Z., Liu, X., Chen, W., Xie, M., Wang, W., Hammond, S., Andersen, M.R., Neff, N., Passarelli, B., Koh, W., Fan, H.C., Wang, J., Gui, Y., Lee, K.H., Betenbaugh, M.J., Quake, S.R., Famili, I., Palsson, B.O., Wang, J., 2011. The genomic sequence of the Chinese hamster ovary (CHO)-K1 cell line. Nature Biotechnology 29, 735–741.

[25] Davis, S. J., & van der Merwe, P. A. 1996. The structure and ligand interactions of CD2: implications for T-cell function. Immunology today, 17(4), 177–187.

[26] Binder C, Cvetkovski F, Sellberg F, Berg S, Paternina Visbal H, Sachs DH, Berglund E, Berglund D. CD2 Immunobiology. 2020. Front Immunol. 11, 1090.

[27] Walker, A.J., Majzner, R.G., Zhang, L., Wanhainen, K., Long, A.H., Nguyen, S.M., Lopomo, P., Vigny, M., Fry, T.J., Orentas, R.J., Mackall, C.L., 2017. Tumor Antigen and Receptor Densities Regulate Efficacy of a Chimeric Antigen Receptor Targeting Anaplastic Lymphoma Kinase. Molecular Therapy 25, 2189–2201.

[28] Purbhoo, M.A., Irvine, D.J., Huppa, J.B., Davis, M.M., 2004. T cell killing does not require the formation of a stable mature immunological synapse. Nature Immunology 5, 524–530.

[29] Huang, J., Brameshuber, M., Zeng, X., Xie, J., Li, Q.-J., Chien, Y.-H., Valitutti, S., Mark, 2013. A Single Peptide-Major Histocompatibility Complex Ligand Triggers Digital Cytokine Secretion in CD4+ T Cells. Immunity 39, 846–857.

[30] Watanabe, K., Terakura, S., Martens, A.C., Van Meerten, T., Uchiyama, S., Imai, M., Sakemura, R., Goto, T., Hanajiri, R., Imahashi, N., Shimada, K., Tomita, A., Kiyoi, H., Nishida, T., Naoe, T., Murata, M., 2015. Target Antigen Density Governs the Efficacy of Anti–CD20-CD28-CD3*ζ* Chimeric Antigen Receptor–Modified Effector CD8+ T Cells. The Journal of Immunology 194, 911–920.

[31] Fry, T.J., Shah, N.N., Orentas, R.J., Stetler-Stevenson, M., Yuan, C.M., Ramakrishna, S., Wolters, P., Martin, S., Delbrook, C., Yates, B., Shalabi, H., Fountaine, T.J., Shern, J.F., Majzner, R.G., Stroncek, D.F., Sabatino, M., Feng, Y., Dimitrov, D.S., Zhang, L., Nguyen, S., Qin, H., Dropulic, B., Lee, D.W., Mackall, C.L., 2018. CD22-targeted CAR T cells induce remission in B-ALL that is naive or resistant to CD19-targeted CAR immunotherapy. Nature Medicine 24, 20–28.

[32] Majzner, R.G., Rietberg, S.P., Sotillo, E., Dong, R., Vachharajani, V.T., Labanieh, L., Myklebust, J.H., Kadapakkam, M., Weber, E.W., Tousley, A.M., Richards, R.M., Heitzeneder, S., Nguyen, S.M., Wiebking, V., Theruvath, J., Lynn, R.C., Xu, P., Dunn, A.R., Vale, R.D., Mackall, C.L., 2020. Tuning the Antigen Density Requirement for CAR T-cell Activity. Cancer Discovery 10, 702–723.

[33] Lynn, R.C., Weber, E.W., Sotillo, E., Gennert, D., Xu, P., Good, Z., Anbunathan, H., Lattin, J., Jones, R., Tieu, V., Nagaraja, S., Granja, J., De Bourcy, C.F.A., Majzner, R., Satpathy, A.T., Quake, S.R., Monje, M., Chang, H.Y., Mackall, C.L., 2019. c-Jun overexpression in CAR T cells induces exhaustion resistance. Nature 576, 293–300.

[34] Ziegler, S. F., Ramsdell, F., & Alderson, M. R. 1994. The activation antigen CD69. Stem cells, 12(5), 456–465.

[35] Cibrián, D., Sánchez-Madrid, F., 2017. CD69: from activation marker to metabolic gatekeeper. European Journal of Immunology 47, 946–953.

[36] Green, D.R., Droin, N., Pinkoski, M., 2003. Activation-induced cell death in T cells. Immunological Reviews 193, 70–81.

[37] Singer, M., Wang, C., Cong, L., Marjanovic, N.D., Kowalczyk, M.S., Zhang, H., Nyman, J., Sakuishi, K., Kurtulus, S., Gennert, D., Xia, J., Kwon, J.Y.H., Nevin, J., Herbst, R.H., Yanai, I., Rozenblatt-Rosen, O., Kuchroo, V.K., Regev, A., Anderson, A.C., 2016. A Distinct Gene Module for Dysfunction Uncoupled from Activation in Tumor-Infiltrating T Cells. Cell 166, 1500–1511.

[38] Steuer, R., Kurths, J., Daub, C. O., Weise, J., & Selbig, J. 2002. The mutual information: detecting and evaluating dependencies between variables. Bioinformatics, 18, S231–S240.

[39] Shannon, C. E., & Weaver, W. 1949. The mathematical theory of communication.

[40] Mi, H., Muruganujan, A., Huang, X., Ebert, D., Mills, C., Guo, X., Thomas, P.D., 2019. Protocol Update for large-scale genome and gene function analysis with the PANTHER classification system (v.14.0). Nature Protocols 14, 703–721.

[41] Thomas, P.D., Ebert, D., Muruganujan, A., Mushayahama, T., Albou, L., Mi, H., 2022. PANTHER : Making genome-scale phylogenetics accessible to all. Protein Science 31, 8–22.

[42] Peters, P. J., Geuze, H. J., van der Donk, H. A., & Borst, J. 1990. A new model for lethal hit delivery by cytotoxic T lymphocytes. Immunology today, 11(1), 28–32.

[43] Fu, W., Lei, C., Liu, S., Cui, Y., Wang, C., Qian, K., Li, T., Shen, Y., Fan, X., Lin, F., Ding, M., Pan, M., Ye, X., Yang, Y., Hu, S., 2019. CAR exosomes derived from effector CAR-T cells have potent antitumour effects and low toxicity. Nature Communications 10.

[44] Monecke, T., Dickmanns, A., & Ficner, R. 2009. Structural basis for m7G-cap hypermethylation of small nuclear, small nucleolar and telomerase RNA by the dimethyltransferase TGS1. Nucleic acids research, 37(12), 3865–3877.

[45] Matera, A.G., Wang, Z., 2014. A day in the life of the spliceosome. Nature Reviews Molecular Cell Biology 15, 108–121.

[46] Plaschka, C., Lin, P.-C., Charenton, C., Nagai, K., 2018. Prespliceosome structure provides insights into spliceosome assembly and regulation. Nature 559, 419–422.

[47] Vercellino, I., Sazanov, L.A., 2022. The assembly, regulation and function of the mitochondrial respiratory chain. Nature Reviews Molecular Cell Biology 23, 141–161.

[48] Bengsch, B., Johnson, A.L., Kurachi, M., Odorizzi, P.M., Pauken, K.E., Attanasio, J., Stelekati, E., Mclane, L.M., Paley, M.A., Delgoffe, G.M., Wherry, E.J., 2016. Bioenergetic Insufficiencies Due to Metabolic Alterations Regulated by the Inhibitory Receptor PD-1 Are an Early Driver of CD8 + T Cell Exhaustion. Immunity 45, 358–373.

[49] Scharping, N.E., Menk, A.V., Moreci, R.S., Whetstone, R.D., Dadey, R.E., Watkins, S.C., Ferris, R.L., Delgoffe, G.M., 2016. The Tumor Microenvironment Represses T Cell Mitochondrial Biogenesis to Drive Intratumoral T Cell Metabolic Insufficiency and Dysfunction. Immunity 45, 374–388.

[50] Vardhana, S.A., Hwee, M.A., Berisa, M., Wells, D.K., Yost, K.E., King, B., Smith, M., Herrera, P.S., Chang, H.Y., Satpathy, A.T., Van Den Brink, M.R.M., Cross, J.R., Thompson, C.B., 2020. Impaired mitochondrial oxidative phosphorylation limits the self-renewal of T cells exposed to persistent antigen. Nature Immunology 21, 1022–1033.

[51] Yu, Y.-R., Imrichova, H., Wang, H., Chao, T., Xiao, Z., Gao, M., Rincon-Restrepo, M., Franco, F., Genolet, R., Cheng, W.-C., Jandus, C., Coukos, G., Jiang, Y.-F., Locasale, J.W., Zippelius, A., Liu, P.-S., Tang, L., Bock, C., Vannini, N., Ho, P.-C., 2020. Disturbed mitochondrial dynamics in CD8+ TILs reinforce T cell exhaustion. Nature Immunology 21, 1540–1551.

[52] Scharping, N.E., Rivadeneira, D.B., Menk, A.V., Vignali, P.D.A., Ford, B.R., Rittenhouse, N.L., Peralta, R., Wang, Y., Wang, Y., Depeaux, K., Poholek, A.C., Delgoffe, G.M., 2021. Mitochondrial stress induced by continuous stimulation under hypoxia rapidly drives T cell exhaustion. Nature Immunology 22, 205–215.

[53] Lisci, M., Barton, P.R., Randzavola, L.O., Ma, C.Y., Marchingo, J.M., Cantrell, D.A., Paupe, V., Prudent, J., Stinchcombe, J.C., Griffiths, G.M., 2021. Mitochondrial translation is required for sustained killing by cytotoxic T cells. Science 374.

[54] Glotzer, M., 2009. The 3Ms of central spindle assembly: microtubules, motors and MAPs. Nature Reviews Molecular Cell Biology 10, 9–20.

[55] Mckinley, K.L., Cheeseman, I.M., 2016. The molecular basis for centromere identity and function. Nature Reviews Molecular Cell Biology 17, 16–29.

[56] Curtis, N.L., Ruda, G.F., Brennan, P., Bolanos-Garcia, V.M., 2020. Deregulation of Chromosome Segregation and Cancer. Annual Review of Cancer Biology 4, 257–278.

[57] Johnson, G. L., & Lapadat, R. 2002. Mitogen-activated protein kinase pathways mediated by ERK, JNK, and p38 protein kinases. Science, 298(5600), 1911–1912.

[58] Ashwell, J.D., 2006. The many paths to p38 mitogen-activated protein kinase activation in the immune system. Nature Reviews Immunology 6, 532–540.

[59] Kim, E. H., & Suresh, M. 2013. Role of PI3K/Akt signaling in memory CD8 T cell differentiation. Frontiers in immunology, 4, 20.

[60] Singh, M.D., Ni, M., Sullivan, J.M., Hamerman, J.A., Campbell, D.J., 2018. B cell adaptor for PI3-kinase (BCAP) modulates CD8+ effector and memory T cell differentiation. Journal of Experimental Medicine 215, 2429–2443.

[61] Lesourne, R., Uehara, S., Lee, J., Song, K.-D., Li, L., Pinkhasov, J., Zhang, Y., Weng, N.-P., Wildt, K.F., Wang, L., Bosselut, R., Love, P.E., 2009. Themis, a T cell–specific protein important for late thymocyte development. Nature Immunology 10, 840–847.

[62] Paster, W., Bruger, A.M., Katsch, K., Grégoire, C., Roncagalli, R., Fu, G., Gascoigne, N.R., Nika, K., Cohnen, A., Feller, S.M., Simister, P.C., Molder, K.C., Cordoba, S., Dushek, O., Malissen, B., Acuto, O., 2015. A THEMIS:SHP 1 complex promotes T-cell survival. The EMBO Journal 34, 393–409.

[63] Liu, Y., Cong, Y., Niu, Y., Yuan, Y., Tan, F., Lai, Q., Hu, Y., Hou, B., Li, J., Lin, C., Zheng, H., Dong, J., Tang, J., Chen, Q., Brzostek, J., Zhang, X., Chen, X.L., Wang, H.-R., Gascoigne, N.R.J., Xu, B., Lin, S.-H., Fu, G., 2022. Themis is indispensable for IL-2 and IL-15 signaling in T cells. Science Signaling 15.

[64] Cook, M. E., Bradstreet, T. R., Webber, A. M., Kim, J., Santeford, A., Harris, K. M., Murphy, M. K., Tran, J., Abdalla, N. M., Schwarzkopf, E. A., Greco, S. C., Halabi, C. M., Apte, R. S., Blackshear, P. J., Edelson, B. T. 2022. The ZFP36 family of RNA binding proteins regulates homeostatic and autoreactive T cell responses. Science immunology, 7, eabo0981.

[65] Wold, F. and Ballou, C.E., 1957. Studies on the enzyme enolase: I. Equilibrium studies. Journal of Biological Chemistry, 227, 301–312.

[66] Huppertz, I., Perez-Perri, J. I., Mantas, P., Sekaran, T., Schwarzl, T., Russo, F., Ferring-Appel, D., Koskova, Z., Dimitrova-Paternoga, L., Kafkia, E., Hennig, J., Neveu, P. A., Patil, K., & Hentze, M. W. (2022). Riboregulation of Enolase 1 activity controls glycolysis and embryonic stem cell differentiation. Molecular cell, 82, 2666–2680.

[67] Zheng, C., Zheng, L., Yoo, J.-K., Guo, H., Zhang, Y., Guo, X., Kang, B., Hu, R., Huang, J.Y., Zhang, Q., Liu, Z., Dong, M., Hu, X., Ouyang, W., Peng, J., Zhang, Z., 2017. Landscape of Infiltrating T Cells in Liver Cancer Revealed by Single-Cell Sequencing. Cell 169, 1342–1356.

[68] Collins, S., Wolfraim, L. A., Drake, C. G., Horton, M. R., & Powell, J. D. 2006. Cutting Edge: TCR-induced NAB2 enhances T cell function by coactivating IL-2 transcription. Journal of immunology, 177, 8301–8305.

[69] Milner, J.J., Toma, C., Yu, B., Zhang, K., Omilusik, K., Phan, A.T., Wang, D., Getzler, A.J., Nguyen, T., Crotty, S., Wang, W., Pipkin, M.E., Goldrath, A.W., 2017. Runx3 programs CD8+ T cell residency in non-lymphoid tissues and tumours. Nature 552, 253–257.

[70] Shan, Q., Zeng, Z., Xing, S., Li, F., Hartwig, S.M., Gullicksrud, J.A., Kurup, S.P., Van Braeckel-Budimir, N., Su, Y., Martin, M.D., Varga, S.M., Taniuchi, I., Harty, J.T., Peng, W., Badovinac, V.P., Xue, H.-H., 2017. The transcription factor Runx3 guards cytotoxic CD8+ effector T cells against deviation towards follicular helper T cell lineage. Nature Immunology 18, 931–939.

[71] Wang, D., Diao, H., Getzler, A.J., Rogal, W., Frederick, M.A., Milner, J., Yu, B., Crotty, S., Goldrath, A.W., Pipkin, M.E., 2018. The Transcription Factor Runx3 Establishes Chromatin Accessibility of cis-Regulatory Landscapes that Drive Memory Cytotoxic T Lymphocyte Formation. Immunity 48, 659–674.

[72] Schumann, K., Raju, S.S., Lauber, M., Kolb, S., Shifrut, E., Cortez, J.T., Skartsis, N., Nguyen, V.Q., Woo, J.M., Roth, T.L., Yu, R., Nguyen, M.L.T., Simeonov, D.R., Nguyen, D.N., Targ, S., Gate, R.E., Tang, Q., Bluestone, J.A., Spitzer, M.H., Ye, C.J., Marson, A., 2020. Functional CRISPR dissection of gene networks controlling human regulatory T cell identity. Nature Immunology 21, 1456–1466.

[73] Sukumar, M., Liu, J., Ji, Y., Subramanian, M., Crompton, J.G., Yu, Z., Roychoudhuri, R., Palmer, D.C., Muranski, P., Karoly, E.D., Mohney, R.P., Klebanoff, C.A., Lal, A., Finkel, T., Restifo, N.P., Gattinoni, L., 2013. Inhibiting glycolytic metabolism enhances CD8+ T cell memory and antitumor function. Journal of Clinical Investigation 123, 4479–4488.

[74] Aibar, S., González-Blas, C.B., Moerman, T., Huynh-Thu, V.A., Imrichova, H., Hulselmans, G., Rambow, F., Marine, J.-C., Geurts, P., Aerts, J., Van Den Oord, J., Atak, Z.K., Wouters, J., Aerts, S., 2017. SCENIC: single-cell regulatory network inference and clustering. Nature Methods 14, 1083–1086.

[75] Matthew, Simon, Heather, Ian, Jonathan, Curtis, Cui, G., Ming, Susan, 2014. The Transcription Factor FoxO1 Sustains Expression of the Inhibitory Receptor PD-1 and Survival of Antiviral CD8+ T Cells during Chronic Infection. Immunity 41, 802–814.

[76] Utzschneider, D.T., Gabriel, S.S., Chisanga, D., Gloury, R., Gubser, P.M., Vasanthakumar, A., Shi, W., Kallies, A., 2020. Early precursor T cells establish and propagate T cell exhaustion in chronic infection. Nature Immunology 21, 1256–1266.

[77] Mclane, L.M., Ngiow, S.F., Chen, Z., Attanasio, J., Manne, S., Ruthel, G., Wu, J.E., Staupe, R.P., Xu, W., Amaravadi, R.K., Xu, X., Karakousis, G.C., Mitchell, T.C., Schuchter, L.M., Huang, A.C., Freedman, B.D., Betts, M.R., Wherry, E.J., 2021. Role of nuclear localization in the regulation and function of T-bet and Eomes in exhausted CD8 T cells. Cell Reports 35 109120.

[78] Yang, Z.-F., Drumea, K., Mott, S., Wang, J., Rosmarin, A.G., 2014. GABP Transcription Factor (Nuclear Respiratory Factor 2) Is Required for Mitochondrial Biogenesis. Molecular and Cellular Biology 34, 3194–3201.

[79] Fan, W., & Evans, R. 2015. PPARs and ERRs: molecular mediators of mitochondrial metabolism. Current opinion in cell biology, 33, 49–54.

[80] Pennycook, B.R., Vesela, E., Peripolli, S., Singh, T., Barr, A.R., Bertoli, C., De Bruin, R.A.M., 2020. E2F-dependent transcription determines replication capacity and S phase length. Nature Communications 11.

[81] Wang, Z. G., Delva, L., Gaboli, M., Rivi, R., Giorgio, M., Cordon-Cardo, C., Grosveld, F., & Pandolfi, P. P. 1998. Role of PML in cell growth and the retinoic acid pathway. Science, 279, 1547–1551.

[82] Scarr, R.B., Smith, M.R., Beddall, M., Sharp, P.A., 2000. A Novel 50-Kilodalton Fragment of Host Cell Factor 1 (C1) in G0 Cells. Molecular and Cellular Biology 20, 3568–3575.

[83] Julien, E., 2003. Proteolytic processing is necessary to separate and ensure proper cell growth and cytokinesis functions of HCF-1. The EMBO Journal 22, 2360–2369.

[84] Liu, Z., Luyten, I., Bottomley, M. J., Messias, A. C., Houngninou-Molango, S., Sprangers, R., Zanier, K., Krämer, A., & Sattler, M. 2001. Structural basis for recognition of the intron branch site RNA by splicing factor 1. Science, 294, 1098–1102.

[85] Schubert, M.-L., Schmitt, M., Wang, L., Ramos, C.A., Jordan, K., Müller-Tidow, C., Dreger, P., 2021. Side-effect management of chimeric antigen receptor (CAR) T-cell therapy. Annals of Oncology 32, 34–48.

[86] Miao, L., Zhang, Z., Ren, Z., & Li, Y. (2021). Reactions Related to CAR-T Cell Therapy. Frontiers in immunology, 12, 663201.

[87] Maude, S.L., Laetsch, T.W., Buechner, J., Rives, S., Boyer, M., Bittencourt, H., Bader, P., Verneris, M.R., Stefanski, H.E., Myers, G.D., Qayed, M., De Moerloose, B., Hiramatsu, H., Schlis, K., Davis, K.L., Martin, P.L., Nemecek, E.R., Yanik, G.A., Peters, C., Baruchel, A., Boissel, N., Mechinaud, F., Balduzzi, A., Krueger, J., June, C.H., Levine, B.L., Wood, P., Taran, T., Leung, M., Mueller, K.T., Zhang, Y., Sen, K., Lebwohl, D., Pulsipher, M.A., Grupp, S.A., 2018. Tisagenlecleucel in Children and Young Adults with B-Cell Lymphoblastic Leukemia. New England Journal of Medicine 378, 439–448.

[88] Jhunjhunwala, S., Hammer, C., Delamarre, L., 2021. Antigen presentation in cancer: insights into tumour immunogenicity and immune evasion. Nature Reviews Cancer 21, 298–312.

[89] Weber, E.W., Parker, K.R., Sotillo, E., Lynn, R.C., Anbunathan, H., Lattin, J., Good, Z., Belk, J.A., Daniel, B., Klysz, D., Malipatlolla, M., Xu, P., Bashti, M., Heitzeneder, S., Labanieh, L., Vandris, P., Majzner, R.G., Qi, Y., Sandor, K., Chen, L.-C., Prabhu, S., Gentles, A.J., Wandless, T.J., Satpathy, A.T., Chang, H.Y., Mackall, C.L., 2021. Transient rest restores functionality in exhausted CAR-T cells through epigenetic remodeling. Science 372, eaba1786.

[90] Zebley, C.C., Youngblood, B., 2022. Mechanisms of T cell exhaustion guiding next-generation immunotherapy. Trends in Cancer 8, 726–734.

[91] Jutz, S., Hennig, A., Paster, W., Asrak, O, Dijanovic, D., Kellner, F., Pickl, W.F., Huppa, J.B., Leitner, J., Steinberger, P., 2017. A cellular platform for the evaluation of immune checkpoint molecules. Oncotarget 8, 64892–64906.

[92] Rosskopf, S., Leitner, J., Paster, W., Morton, L.T., Hagedoorn, R.S., Steinberger, P., Heemskerk, M.H.M., 2018. A Jurkat 76 based triple parameter reporter system to evaluate TCR functions and adoptive T cell strategies. Oncotarget 9, 17608–17619.

[93] Abu-Shah, E., Trendel, N., Kruger, P., Nguyen, J., Pettmann, J., Kutuzov, M., Dushek, O., 2020. Human CD8+ T Cells Exhibit a Shared Antigen Threshold for Different Effector Responses. The Journal of Immunology 205, 1503–1512.

[94] Burton, J., Siller-Farfán, J.A., Pettmann, J., Salzer, B., Kutuzov, M., Van Der Merwe, P.A., Dushek, O., 2023. Inefficient exploitation of accessory receptors reduces the sensitivity of chimeric antigen receptors. Proceedings of the National Academy of Sciences 120.

[95] Lajoie M.J., Boyken S.E., Salter A.I., Bruffey J., Rajan A., Langan R.A., Olshefsky A., Muhunthan V., Bick M.J., Gewe M., Quijano-Rubio A., Johnson J., Lenz G., Nguyen A., Pun S., Correnti C.E., Riddell S.R., Baker D., 2020. Designed protein logic to target cells with precise combinations of surface antigens. Science 369, 1637–1643.

[96] Salter A.I., Rajan A., Kennedy J.J., Ivey R.G., Shelby S.A., Leung I., Templeton M.L., Muhunthan V., Voillet V., Sommermeyer D., Whiteaker J.R., Gottardo R., Veatch S.L., Paulovich A.G., Riddell SR., 2021. Comparative analysis of TCR and CAR signaling informs CAR designs with superior antigen sensitivity and in vivo function. Sci Signal. 14, eabe2606.

[97] Satija, R., Farrell, J.A., Gennert, D., Schier, A.F., Regev, A., 2015. Spatial reconstruction of single-cell gene expression data. Nature Biotechnology 33, 495–502.

[98] Lun, A.T.L., Riesenfeld, S., Andrews, T., Dao, T.P., Gomes, T., Marioni, J.C., 2019. Empty-Drops: distinguishing cells from empty droplets in droplet-based single-cell RNA sequencing data. Genome Biology 20.

[99] Germain, P.-L., Lun, A., Garcia Meixide, C., Macnair, W., Robinson, M.D., 2022. Doublet identification in single-cell sequencing data using scDblFinder. F1000Research 10, 979.

[100] Ahern, D.J., Ai, Z., Ainsworth, M., Allan, C., Allcock, A., Angus, B., Ansari, M.A., Arancibia- Cárcamo, C.V., Aschenbrenner, D., Attar, M., Baillie, J.K., Barnes, E., Bashford-Rogers, R., Bashyal, A., Beer, S., Berridge, G., Beveridge, A., Bibi, S., Bicanic, T., Blackwell, L., Bowness, P., Brent, A., Brown, A., Broxholme, J., Buck, D., Burnham, K.L., Byrne, H., Camara, S., Candido Ferreira, I., Charles, P., Chen, W., Chen, Y.-L., Chong, A., Clutterbuck, E.A., Coles, M., Conlon, C.P., Cornall, R., Cribbs, A.P., Curion, F., Davenport, E.E., Davidson, N., Davis, S., Dendrou, C.A., Dequaire, J., Dib, L., Docker, J., Dold, C., Dong, T., Downes, D., Drakesmith, H., Dunachie, S.J., Duncan, D.A., Eijsbouts, C., Esnouf, R., Espinosa, A., Etherington, R., Fairfax, B., Fairhead, R., Fang, H., Fassih, S., Felle, S., Fernandez Mendoza, M., Ferreira, R., Fischer, R., Foord, T., Forrow, A., Frater, J., Fries, A., Gallardo Sanchez, V., Garner, L.C., Geeves, C., Georgiou, D., Godfrey, L., Golubchik, T., Gomez Vazquez, M., Green, A., Harper, H., Harrington, H.A., Heilig, R., Hester, S., Hill, J., Hinds, C., Hird, C., Ho, L.-P., Hoekzema, R., Hollis, B., Hughes, J., Hutton, P., Jackson-Wood, M.A., Jainarayanan, A., James-Bott, A., Jansen, K., Jeffery, K., Jones, E., Jostins, L., Kerr, G., Kim, D., Klenerman, P., Knight, J.C., Kumar, V., Kumar Sharma, P., Kurupati, P., Kwok, A., Lee, A., Linder, A., Lockett, T., Lonie, L., Lopopolo, M., Lukoseviciute, M., Luo, J., Marinou, S., Marsden, B., Martinez, J., Matthews, P.C., Mazurczyk, M., Mcgowan, S., Mckechnie, S., Mead, A., Mentzer, A.J., Mi, Y., Monaco, C., Montadon, R., Napolitani, G., Nassiri, I., Novak, A., O’Brien, D.P., O’Connor, D., O’Donnell, D., Ogg, G., Overend, L., Park, I., Pavord, I., Peng, Y., Penkava, F., Pereira Pinho, M., Perez, E., Pollard, A.J., Powrie, F., Psaila, B., Quan, T.P., Repapi, E., Revale, S., Silva-Reyes, L., Richard, J.-B., Rich-Griffin, C., Ritter, T., Rollier, C.S., Rowland, M., Ruehle, F., Salio, M., Sansom, S.N., Sanches Peres, R., Santos Delgado, A., Sauka-Spengler, T., Schwessinger, R., Scozzafava, G., Screaton, G., Seigal, A., Semple, M.G., Sergeant, M., Simoglou Karali, C., Sims, D., Skelly, D., Slawinski, H., Sobrinodiaz, A., Sousos, N., Stafford, L., Stockdale, L., Strickland, M., Sumray, O., Sun, B., Taylor, C., Taylor, S., Taylor, A., Thongjuea, S., Thraves, H., Todd, J.A., Tomic, A., Tong, O., Trebes, A., Trzupek, D., Tucci, F.A., Turtle, L., Udalova, I., Uhlig, H., Van Grinsven, E., Vendrell, I., Verheul, M., Voda, A., Wang, G., Wang, L., Wang, D., Watkinson, P., Watson, R., Weinberger, M., Whalley, J., Witty, L., Wray, K., Xue, L., Yeung, H.Y., Yin, Z., Young, R.K., Youngs, J., Zhang, P., Zurke, Y.-X., 2022. A blood atlas of COVID-19 defines hallmarks of disease severity and specificity. Cell 185, 916–938.e58.

[101] Dobin A, Davis CA, Schlesinger F, Drenkow J, Zaleski C, Jha S, Batut P, Chaisson M, Gingeras TR., 2013. STAR: ultrafast universal RNA-seq aligner. Bioinformatics. 29, 15–21.

[102] Huang, X., Huang, Y., 2021. Cellsnp-lite: an efficient tool for genotyping single cells. Bioinformatics 37, 4569–4571.

[103] Huang, Y., Mccarthy, D.J., Stegle, O., 2019. Vireo: Bayesian demultiplexing of pooled single-cell RNA-seq data without genotype reference. Genome Biology 20.

[104] Morgan M., Anders S., Lawrence M., Aboyoun P., Pagès H., Gentleman R., 2009. ShortRead: a bioconductor package for input, quality assessment and exploration of high-throughput sequence data. Bioinformatics 25, 2607–8.

[105] Andreatta, M., Carmona, S.J., 2021. UCell: Robust and scalable single-cell gene signature scoring. Computational and Structural Biotechnology Journal 19, 3796–3798.

[106] Mi, H., Muruganujan, A., Huang, X., Ebert, D., Mills, C., Guo, X., Thomas, P.D., 2019. Protocol Update for large-scale genome and gene function analysis with the PANTHER classification system (v.14.0). Nature Protocols 14, 703–721.

[107] Langfelder, P., Horvath, S., 2008. WGCNA: an R package for weighted correlation network analysis. BMC Bioinformatics 9, 559.

[108] Ali, S., Kjeken, R., Niederlaender, C., Markey, G., Saunders, T.S., Opsata, M., Moltu, K., Bremnes, B., Grønevik, E., Muusse, M., Håkonsen, G.D., Skibeli, V., Kalland, M.E., Wang, I., Buajordet, I., Urbaniak, A., Johnston, J., Rantell, K., Kerwash, E., Schuessler-Lenz, M., Salmonson, T., Bergh, J., Gisselbrecht, C., Tzogani, K., Papadouli, I., Pignatti, F., 2020. The European Medicines Agency Review of Kymriah (Tisagenlecleucel) for the Treatment of Acute Lymphoblastic Leukemia and Diffuse Large B-Cell Lymphoma. The Oncologist 25, e321–e327.

[109] Chan, W.K., Suwannasaen, D., Throm, R.E., Li, Y., Eldridge, P.W., Houston, J., Gray, J.T., Pui, C.-H., Leung, W., 2015. Chimeric antigen receptor-redirected CD45RA-negative T cells have potent antileukemia and pathogen memory response without graft-versus-host activity. Leukemia 29, 387–395.

[110] Mcgeary, S.E., Lin, K.S., Shi, C.Y., Pham, T.M., Bisaria, N., Kelley, G.M., Bartel, D.P., 2019. The biochemical basis of microRNA targeting efficacy. Science 366, eaav1741.

[111] Gargett, T., Yu, W., Dotti, G., Yvon, E.S., Christo, S.N., Hayball, J.D., Lewis, I.D., Brenner, M.K., Brown, M.P., 2016. GD2-specific CAR T Cells Undergo Potent Activation and Deletion Following Antigen Encounter but can be Protected From Activation-induced Cell Death by PD-1 Blockade. Molecular Therapy 24, 1135–1149.

[112] Wherry, E. J., & Kurachi, M. 2015. Molecular and cellular insights into T cell exhaustion. Nature Reviews Immunology, 15, 486–499.

[113] Kawalekar, O.U., O’Connor, R.S., Fraietta, J.A., Guo, L., Mcgettigan, S.E., Posey, A.D., Patel, P.R., Guedan, S., Scholler, J., Keith, B., Snyder, N.W., Blair, I.A., Milone, M.C., June, C.H., 2016. Distinct Signaling of Coreceptors Regulates Specific Metabolism Pathways and Impacts Memory Development in CAR T Cells. Immunity 44, 380–390.

[114] Selli, M. E., Landmann, J. H., Terekhova, M., Lattin, J., Heard, A., Hsu, Y. S., Chang, T., C., Chang, J., Warrington, J., Ha, H., Kingston, N., Hogg, G., Slade, M., Berrien-Elliott, M. M., Foster, M., Kersting-Schadek, S., Gruszczynska, A., DeNardo, D., Fehniger, T. A., Artyomov, M., Singh, N. (2023). Costimulatory domains direct distinct fates of CAR-driven T-cell dysfunction. Blood, 141, 3153–3165.

[115] Venteicher, A.S., Abreu, E.B., Meng, Z., Mccann, K.E., Terns, R.M., Veenstra, T.D., Terns, M.P., Artandi, S.E., 2009. A Human Telomerase Holoenzyme Protein Required for Cajal Body Localization and Telomere Synthesis. Science 323, 644–648.

[116] Albelda, S.M., 2024. CAR T cell therapy for patients with solid tumours: key lessons to learn and unlearn. Nature Reviews Clinical Oncology 21, 47–66.

[117] Zebley, C.C., Zehn, D., Gottschalk, S., Chi, H., 2024. T cell dysfunction and therapeutic intervention in cancer. Nature Immunology 25, 1344–1354.

[118] Brudno, J.N., Kochenderfer, J.N., 2024. Current understanding and management of CAR T cell-associated toxicities. Nature Reviews Clinical Oncology 21, 501–521.

[119] Cordas Dos Santos, D.M., Tix, T., Shouval, R., Gafter-Gvili, A., Alberge, J.-B., Cliff, E.R.S., Theurich, S., Von Bergwelt-Baildon, M., Ghobrial, I.M., Subklewe, M., Perales, M.-A., Rejeski, K., 2024. A systematic review and meta-analysis of nonrelapse mortality after CAR T cell therapy. Nature Medicine 30, 2667–2678.

[120] Cappell, K.M., Kochenderfer, J.N., 2023. Long-term outcomes following CAR T cell therapy: what we know so far. Nature Reviews Clinical Oncology 20, 359–371.

[121] Davila, M.L., Brentjens, R.J., 2022. CAR T cell therapy: looking back and looking forward. Nature Cancer 3, 1418–1419.

[122] Melenhorst, J.J., Chen, G.M., Wang, M., Porter, D.L., Chen, C., Collins, M.A., Gao, P., Bandyopadhyay, S., Sun, H., Zhao, Z., Lundh, S., Pruteanu-Malinici, I., Nobles, C.L., Maji, S., Frey, N.V., Gill, S.I., Loren, A.W., Tian, L., Kulikovskaya, I., Gupta, M., Ambrose, D.E., Davis, M. M., Fraietta, J.A., Brogdon, J.L., Young, R.M., Chew, A., Levine, B.L., Siegel, D.L., Alanio, C., Wherry, E.J., Bushman, F.D., Lacey, S.F., Tan, K., June, C.H., 2022. Decade-long leukaemia remissions with persistence of CD4+ CAR T cells. Nature 602, 503–509.

[123] Morris, E.C., Neelapu, S.S., Giavridis, T., Sadelain, M., 2022. Cytokine release syndrome and associated neurotoxicity in cancer immunotherapy. Nature Reviews Immunology 22, 85–96.

[124] Vinnakota, J.M., Biavasco, F., Schwabenland, M., Chhatbar, C., Adams, R.C., Erny, D., Duquesne, S., El Khawanky, N., Schmidt, D., Fetsch, V., Zähringer, A., Salié, H., Athanassopoulos, D., Braun, L.M., Javorniczky, N.R., Ho, J.N.H.G., Kierdorf, K., Marks, R., Wäsch, R., Simonetta, F., Andrieux, G., Pfeifer, D., Monaco, G., Capitini, C., Fry, T.J., Blank, T., Blazar, B.R., Wagner, E., Theobald, M., Sommer, C., Stelljes, M., Reicherts, C., Jeibmann, A., Schittenhelm, J., Monoranu, C.-M., Rosenwald, A., Kortüm, M., Rasche, L., Einsele, H., Meyer, P.T., Brumberg, J., Völkl, S., Mackensen, A., Coras, R., Von Bergwelt-Baildon, M., Albert, N.L., Bartos, L.M., Brendel, M., Holzgreve, A., Mack, M., Boerries, M., Mackall, C.L., Duyster, J., Henneke, P., Priller, J., Köhler, N., Strübing, F., Bengsch, B., Ruella, M., Subklewe, M., Von Baumgarten, L., Gill, S., Prinz, M., Zeiser, R., 2024. Targeting TGF-activated kinase-1 activation in microglia reduces CAR T immune effector cell-associated neurotoxicity syndrome. Nature Cancer 5, 1227–1249.

[125] Neelapu, S.S., Locke, F.L., Bartlett, N.L., Lekakis, L.J., Miklos, D.B., Jacobson, C.A., Braunschweig, I., Oluwole, O.O., Siddiqi, T., Lin, Y., Timmerman, J.M., Stiff, P.J., Friedberg, J.W., Flinn, I.W., Goy, A., Hill, B.T., Smith, M.R., Deol, A., Farooq, U., Mcsweeney, P., Munoz, J., Avivi, I., Castro, J.E., Westin, J.R., Chavez, J.C., Ghobadi, A., Komanduri, K.V., Levy, R., Jacobsen, E.D., Witzig, T.E., Reagan, P., Bot, A., Rossi, J., Navale, L., Jiang, Y., Aycock, J., Elias, M., Chang, D., Wiezorek, J., Go, W.Y., 2017. Axicabtagene Ciloleucel CAR T-Cell Therapy in Refractory Large B-Cell Lymphoma. New England Journal of Medicine 377, 2531–2544.

[126] Locke, F.L., Neelapu, S.S., Bartlett, N.L., Siddiqi, T., Chavez, J.C., Hosing, C.M., Ghobadi, A., Budde, L.E., Bot, A., Rossi, J.M., Jiang, Y., Xue, A.X., Elias, M., Aycock, J., Wiezorek, J., Go, W.Y., 2017. Phase 1 Results of ZUMA-1: A Multicenter Study of KTE-C19 Anti-CD19 CAR T Cell Therapy in Refractory Aggressive Lymphoma. Molecular Therapy 25, 285–295.

[127] Reina-Campos, M., Scharping, N.E., Goldrath, A.W., 2021. CD8+ T cell metabolism in infection and cancer. Nature Reviews Immunology 21, 718–738.

[128] Keating, S. J., Gu, T., Jun, M. P., & McBride, A. 2022. Health Care Resource Utilization and Total Costs of Care Among Patients with Diffuse Large B Cell Lymphoma Treated with Chimeric Antigen Receptor T Cell Therapy in the United States. Transplantation and cellular therapy, 28(7), 404.e1–404.e6.

[129] Badaracco, J., Gitlin, M., & Keating, S. J. 2023. A Model to Estimate Cytokine Release Syndrome and Neurological Event Management Costs Associated With CAR T-Cell Therapy. Transplantation and cellular therapy, 29(1), 59.e1–59.e6.

[130] Cummings Joyner, A.K., Snider, J.T., Wade, S.W., Wang, S.-T., Buessing, M.G., Johnson, S., Gergis, U., 2022. Cost-Effectiveness of Chimeric Antigen Receptor T Cell Therapy in Patients with Relapsed or Refractory Large B Cell Lymphoma: No Impact of Site of Care. Advances in Therapy 39, 3560–3577.

[131] Abramson, J.S., Siddiqi, T., Garcia, J., Dehner, C., Kim, Y., Nguyen, A., Snyder, S., Mcgarvey, N., Gitlin, M., Pelletier, C., Jun, M.P., 2021. Cytokine release syndrome and neurological event costs in lisocabtagene maraleucel–treated patients in the TRANSCEND NHL 001 trial. Blood Advances 5, 1695–1705.

[132] Lyman, G.H., Nguyen, A., Snyder, S., Gitlin, M., Chung, K.C., 2020. Economic Evaluation of Chimeric Antigen Receptor T-Cell Therapy by Site of Care Among Patients With Relapsed or Refractory Large B-Cell Lymphoma. JAMA Network Open 3, e202072.

[133] Maziarz, R.T., Yang, H., Liu, Q., Wang, T., Zhao, J., Lim, S., Lee, S., Dalal, A., Bollu, V., 2022. Real-world healthcare resource utilization and costs associated with tisagenlecleucel and axicabtagene ciloleucel among patients with diffuse large B-cell lymphoma: an analysis of hospital data in the United States. Leukemia & Lymphoma 63, 2052–2062.

[134] Zhou, X., Cao, H., Fang, S.-Y., Chow, R.D., Tang, K., Majety, M., Bai, M., Dong, M.B., Renauer, P.A., Shang, X., Suzuki, K., Levchenko, A., Chen, S., 2023. CTLA-4 tail fusion enhances CAR-T antitumor immunity. Nature Immunology 24, 1499–1510.

