## Extended data figures 1-17 for "Precise engineering of chimeric antigen receptor expression levels defines T cell identity and function"

[1]MRC Weatherall Institute of Molecular Medicine, Radcliffe Department of Medicine, University of Oxford, Oxford, UK.
[2]Fred Hutchinson Cancer Center, University of Washington, 1100 Fairview Ave N, Seattle, Washington, USA.
[3]Sir William Dunn School of Pathology, University of Oxford, S Parks Rd, Oxford, UK.
[4]MRC Centre for Computational Biology, MRC Weatherall Institute of Molecular Medicine, University of Oxford, Oxford, UK.
[5]Paul Albrechtsen Research Institute, CancerCare Manitoba, Winnipeg, Canada.
[6]Department of Biochemistry & Medical Genetics, Rady Faculty of Health Sciences, University of Manitoba, Winnipeg, Manitoba, Canada.
[7]Stowers Institute for Medical Research, 1000 E 50 St, Kansas City, Missouri, USA.
[†]Present address: Department of Biochemistry, University of Zurich, Winterthurerstrasse 190, Zurich, Switzerland.
[§]Present address: Vertex Pharmaceuticals, 50 Northern Avenue, Boston, MA, USA.

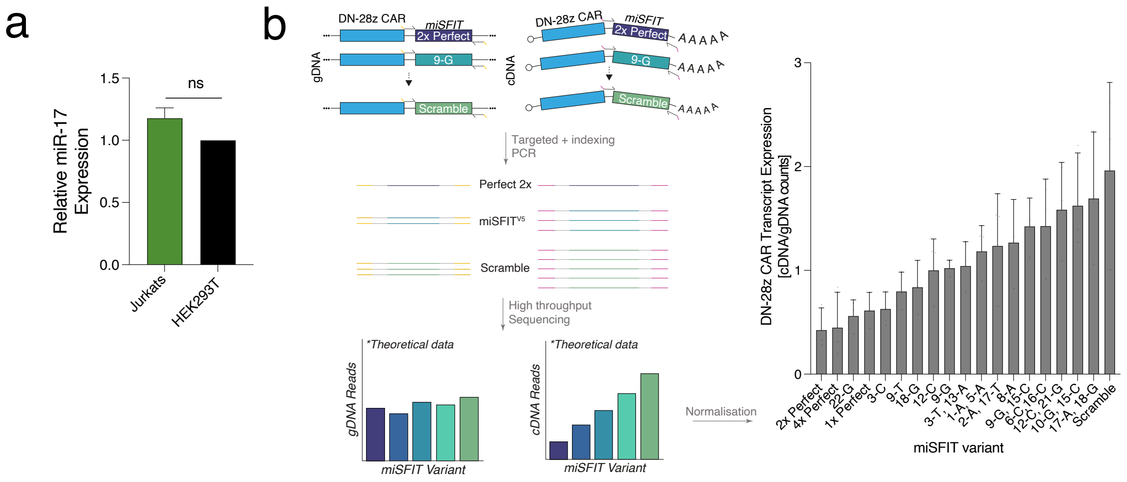


**Extended Data Figure 1**: Establishing a model of tuned CAR expression. **a)** RT-qPCR of relative miR-17 expression in HEK293T and Jurkat T cell line. *RNU6b* miRNA expression was used as a reference for normalisation using the $\Delta\Delta$CT method. Error bars depict standard deviation. N = 3 biological replicates. **b)** Using high-throughput sequencing to measure tuning of the DN-28z CAR transcript. Schematic depicting strategy to measure DN-28z CAR transcript expression. Average, normalised DN-28z CAR transcript expression, calculated by dividing respective miSFIT's total RNA counts (cDNA) divided by total DNA counts (gDNA). N=3 biological replicates, error bars depict standard deviation.

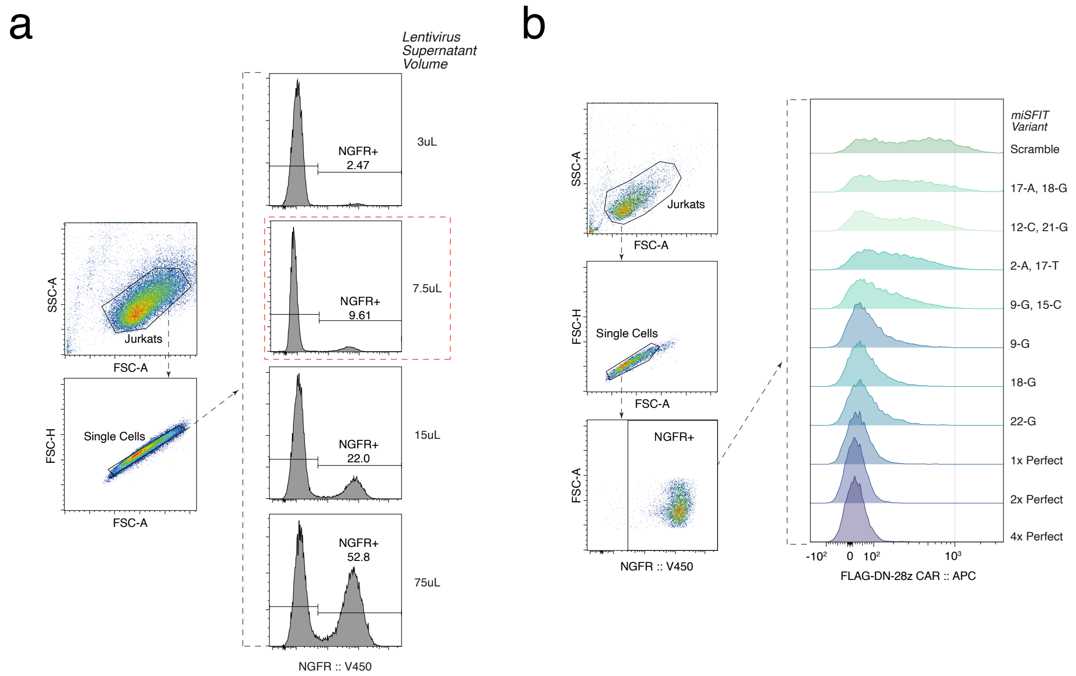


**Extended Data Figure 2**: miSFIT tune CAR surface expression in Jurkat T cells. **a)** Titration of lentivirus volume used to transduce Jurkat T cells for multiplicity of infection (MOI) < 0.1 to ensure single integrands. Selected lentivirus volume boxed in red, used for generation of cell lines shown in **(b)**. Gating strategy and flow cytometry plots of transduction rate, measured by NGFR. **b)** Representative gating strategy and flow cytometry plots of DN-28z CAR expression, measured by staining for FLAG epitope, incorporated within CAR extracellular domain, and flow cytometry.

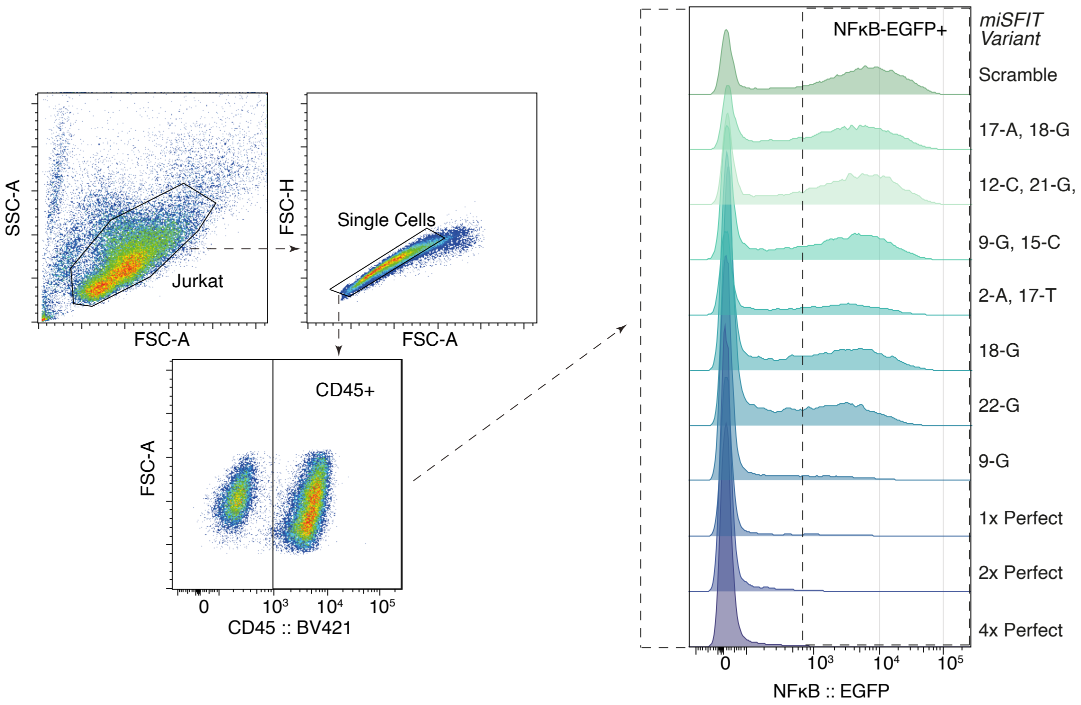


**Extended Data Figure 3**: Tuned CAR expression tunes T cell activation. We co-cultured the DN-28z CAR miSFIT cell lines with synthetic target cells, pulsed with serial dilutions of NY-ESO-1 peptide. Each DN-28z CAR miSFIT cell line contained a reporter of NF-$\kappa$B activity that produced EGFP as readout (**Figure 1c-e**). Representative gating strategy for CAR-miSFIT T cells and flow cytometry plots of EGFP readout for single concentration [20nM] of NY-ESO-1.

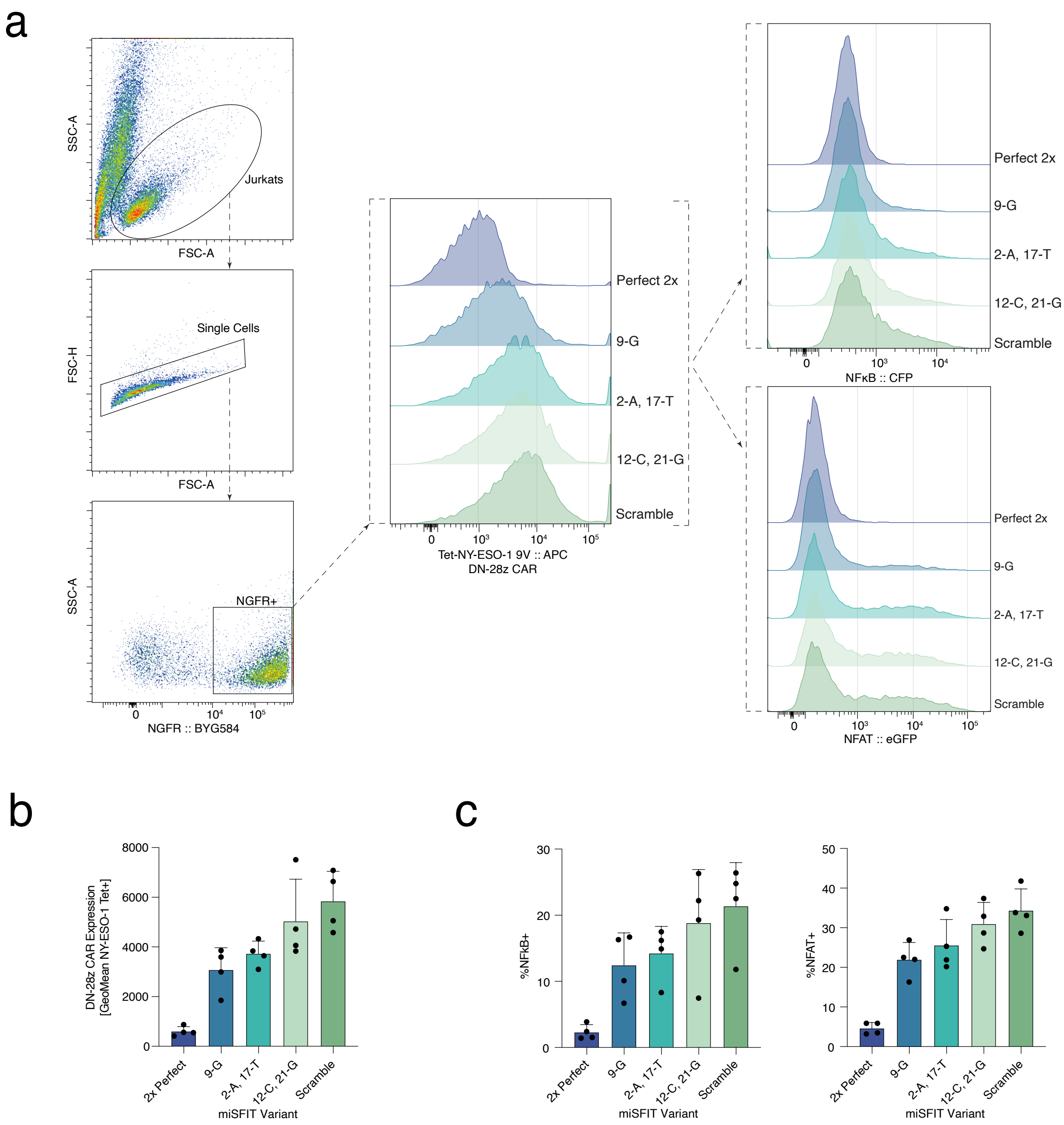


**Extended Data Figure 4**: CAR expression has a direct role in tonic signalling. We introduced a panel of DN-28z CAR miSFIT into a Jurkat cell line with a reporter for NF-$\kappa$B and NFAT and measured basal reporter activity by flow cytometry. **a)** Representative gating strategy and flow cytometry plots of DN-28z CAR, NF-$\kappa$B, and NFAT expression. **b-c)** Average expression of DN-28z CAR, NF-$\kappa$B, and NFAT expression. N=4 biological replicates. Error bars depict standard deviation.

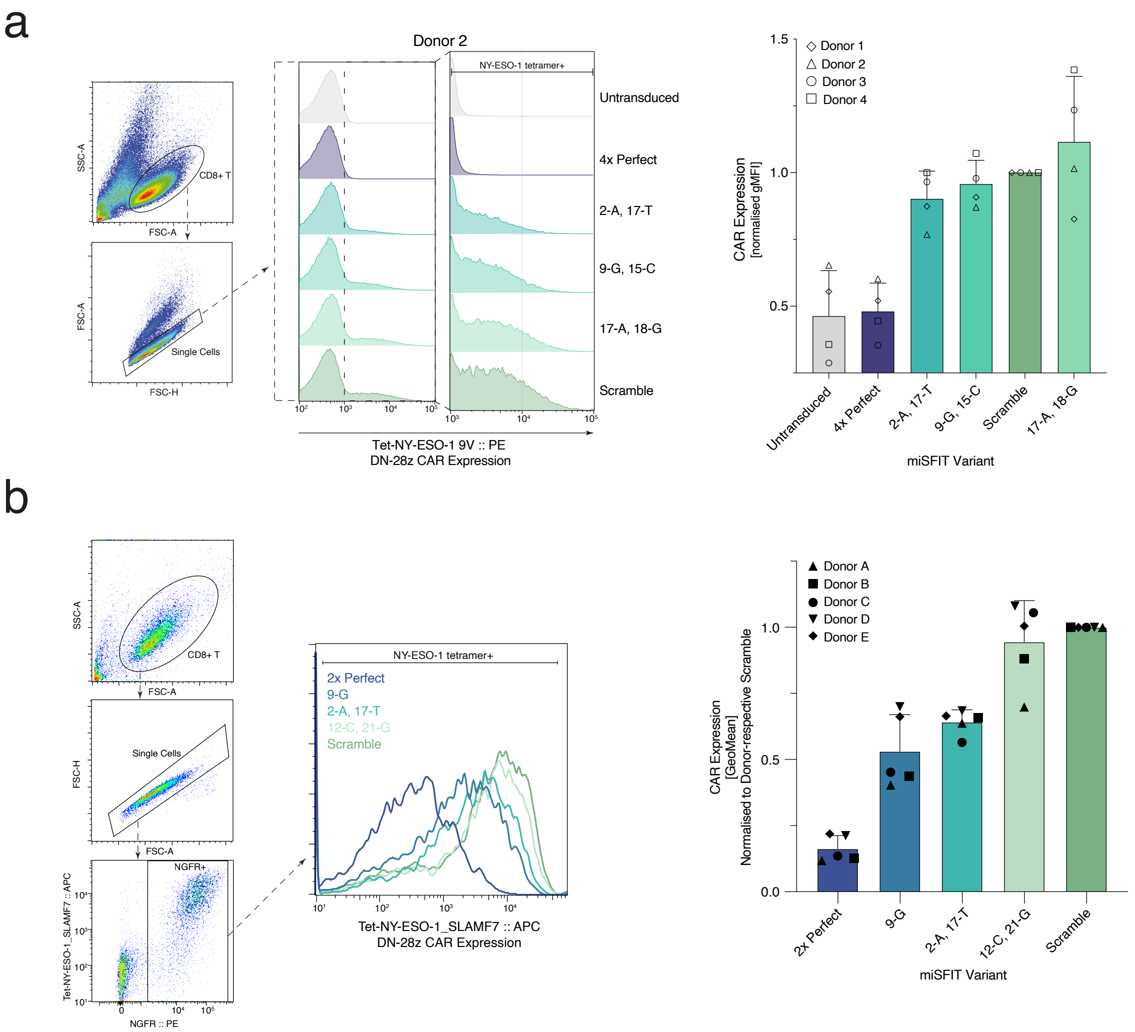


**Extended Data Figure 5**: miSFIT tune CAR expression in primary T cells. Two panels **a & b)** of DN-28z CAR miSFIT were transduced into human, primary CD8+ T cells and average expression was measured by staining with NY-ESO-1 tetramer and flow cytometry. Representative gating strategy and flow cytometry plots and average of DN-28z CAR expression within human primary CD8+ T cells. CAR expression was normalised on a donor basis by dividing the expression of samples by respective Scramble control. **a)** N=4 biological replicates and **b)** N=5 biological replicates. Error bars depict standard deviation.

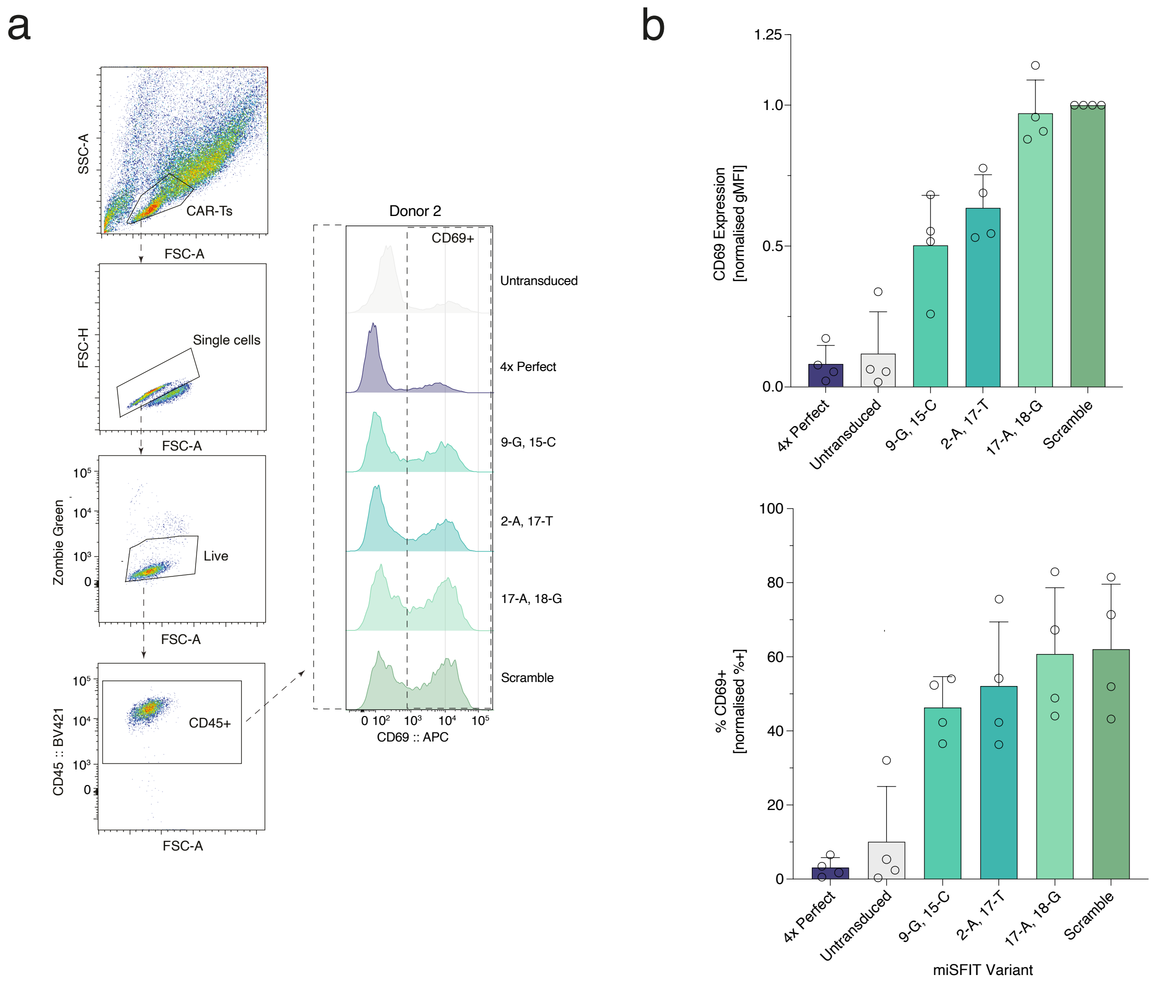


**Extended Data Figure 6**: Tuned CAR expression controls T cell activation in primary T cells. We co-cultured a panel of DN-28z CAR miSFIT T cells with NY-ESO-1-presenting target cells and measured CD69 expression by flow cytometry to verify CAR expression's role on T cell activation. **a)** Representative gating strategy and flow cytometry plots of CD69 expression. **b)** Average MFI and %+ of CD69 expression per miSFIT cell line, normalised on a donor basis by dividing the expression of samples by donor-respective Scramble control. N=4 biological replicates, 2:1 Effector:Target ratio, 20$\mu$M NY-ESO-1 9V peptide concentration. Error bars depict standard deviation.

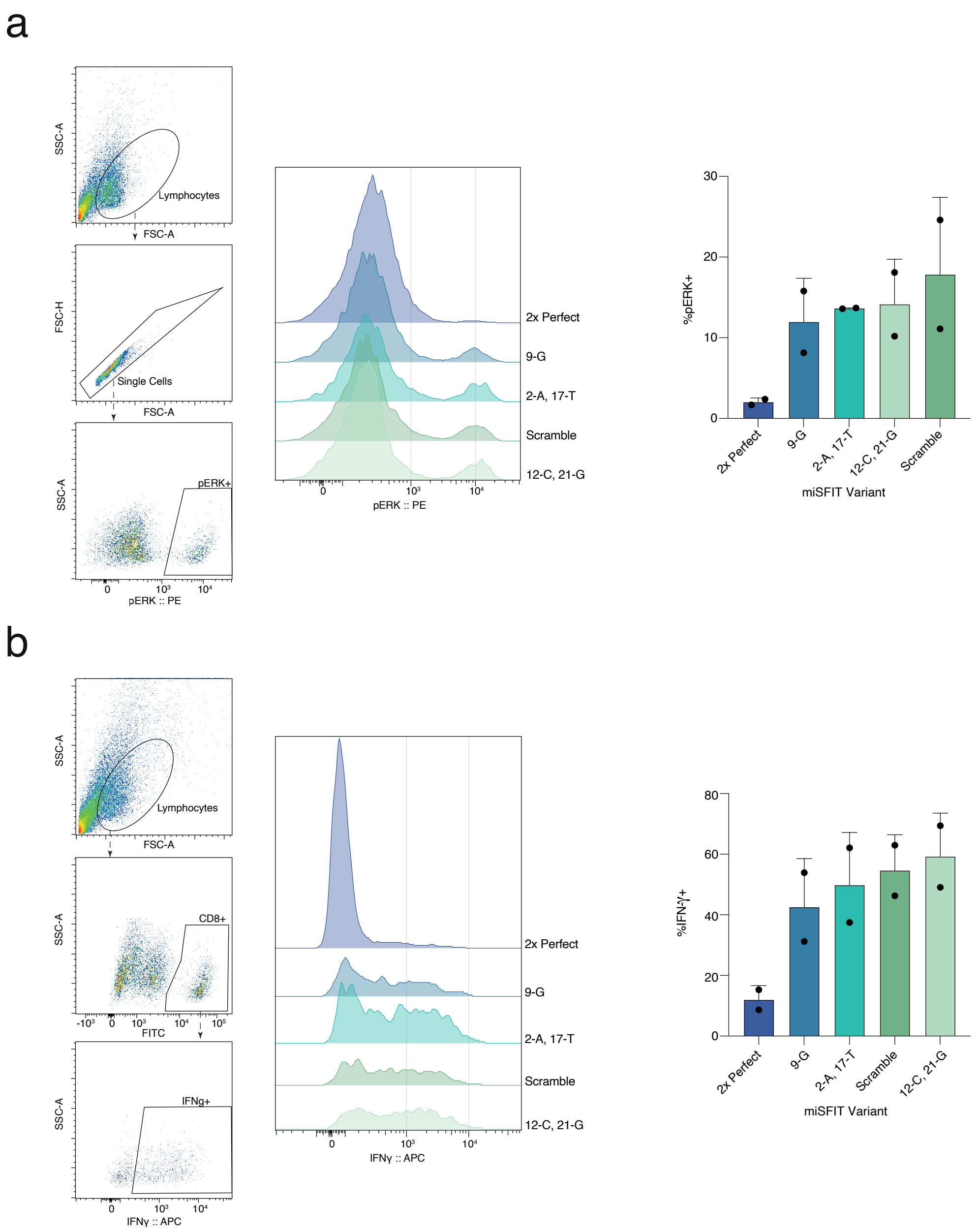


**Extended Data Figure 7**: Tuning CAR expression influences T cell sensitivity. To assay CAR expression's influence on T cell signalling and cytokine production, we exposed a panel of DN-28z CAR miSFIT T cells with titrations of NY-ESO-1 peptide and used flow cytometry to measure **a)** phosphorylation of extracellular signal-regulated kinases (pERK) and **b)** interferon-$\gamma$ production via intracellular cytokine staining. Representative gating strategy, flow cytometry plots, and average % of **a)** phosphorylated ERK+ and **b)** Interferon-$\gamma$+ cells by CAR-miSFIT cell line at 1.25 ug/mL NY-ESO-1 9V peptide concentration. N=2 biological replicates. Error bars depict standard deviation.

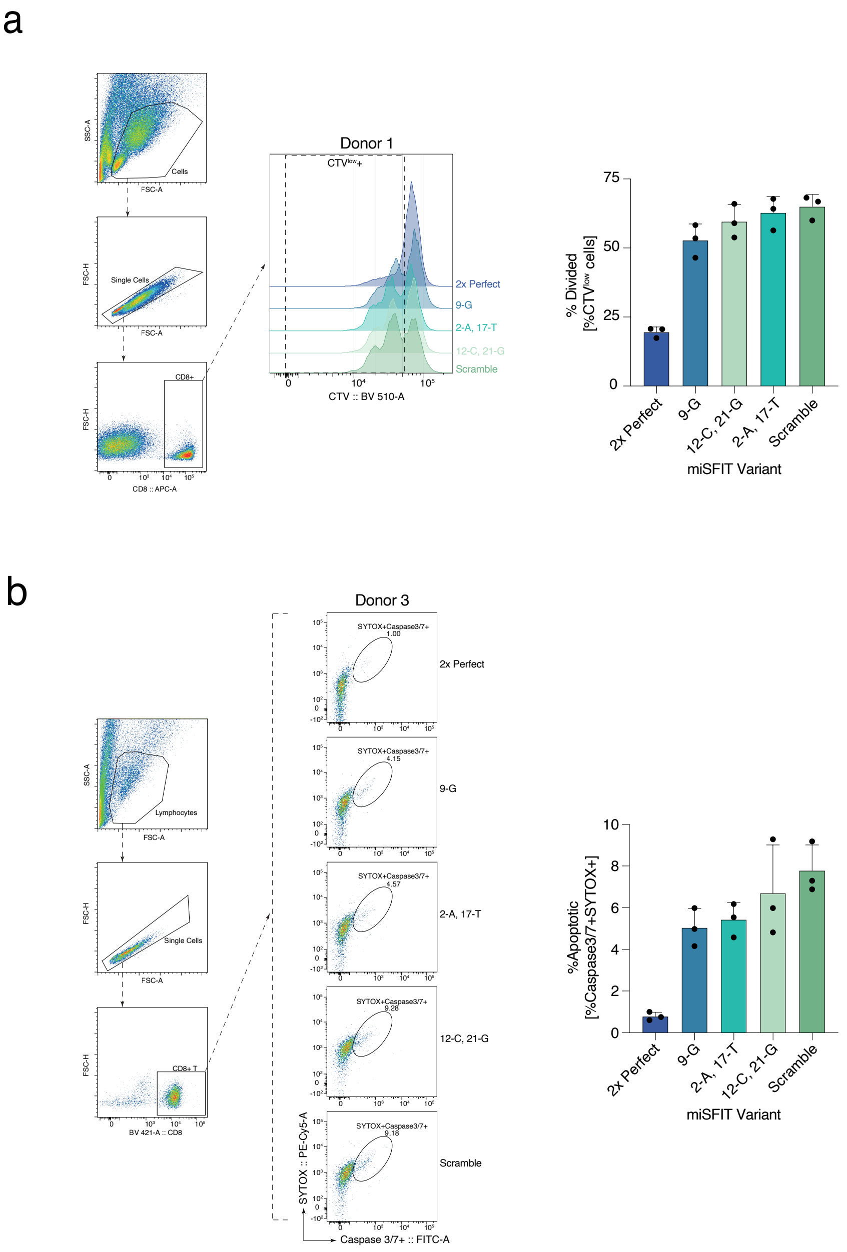


**Extended Data Figure 8**: Tuned CAR expression determines T cell proliferation and activation induced cell-death (AICD) rate. We co-cultured a panel of DN-28z CAR miSFIT T cells with NY-ESO-1 presenting target cells and **a)** used CellTrace Violet (CTV, Methods and Materials) to assess T cell proliferation and **b)** stained for Caspase-3/7 and SYTOX (Methods and Materials) to measure AICD. **a)** Gating strategy, flow cytometry plots, and average %Divided per CAR-miSFIT T cell line. 1:2 Effector:Target ratio. N=3 biological replicates, 0.01$\mu$M NY-ESO-1 9V peptide concentration. **b)** Gating strategy, representative flow cytometry plots, and average %SYTOX+Caspase3/7+ cells per CAR-miSFIT T cell line. 1:2 Effector: Target ratio. N=3 biological replicates, 0.1$\mu$M NY-ESO-1 9V peptide concentration. Error bars depict standard deviation.

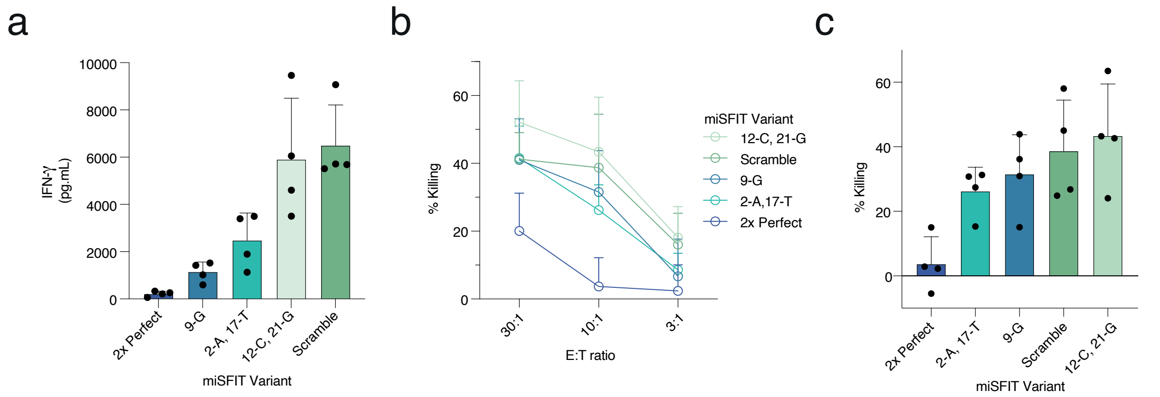


**Extended Data Figure 9**: CAR expression governs effector T cell function. To explore if CAR expression can fine tune effector function, we co-cultured a selection of DN-28z CAR miSFIT T cells with NY-ESO-1 presenting target cells and measured **a)** cytokine secretion and **b,c)** targeted cell lysis. Average **a)** IFN-$\gamma$ secretion and **c)** cytotoxicity (% Killing by $^{51}$Cr release) per CAR-miSFIT T cell line. **b)** Average cytotoxicity by E:T ratio. **a)** 1:1 effector: target ratio and **c)** cytotoxicity 10:1 E:T. **a-c**) N=4 biological replicates, 10$\mu$M NY-ESO-1 9V peptide concentration, and error bars depict standard deviation.

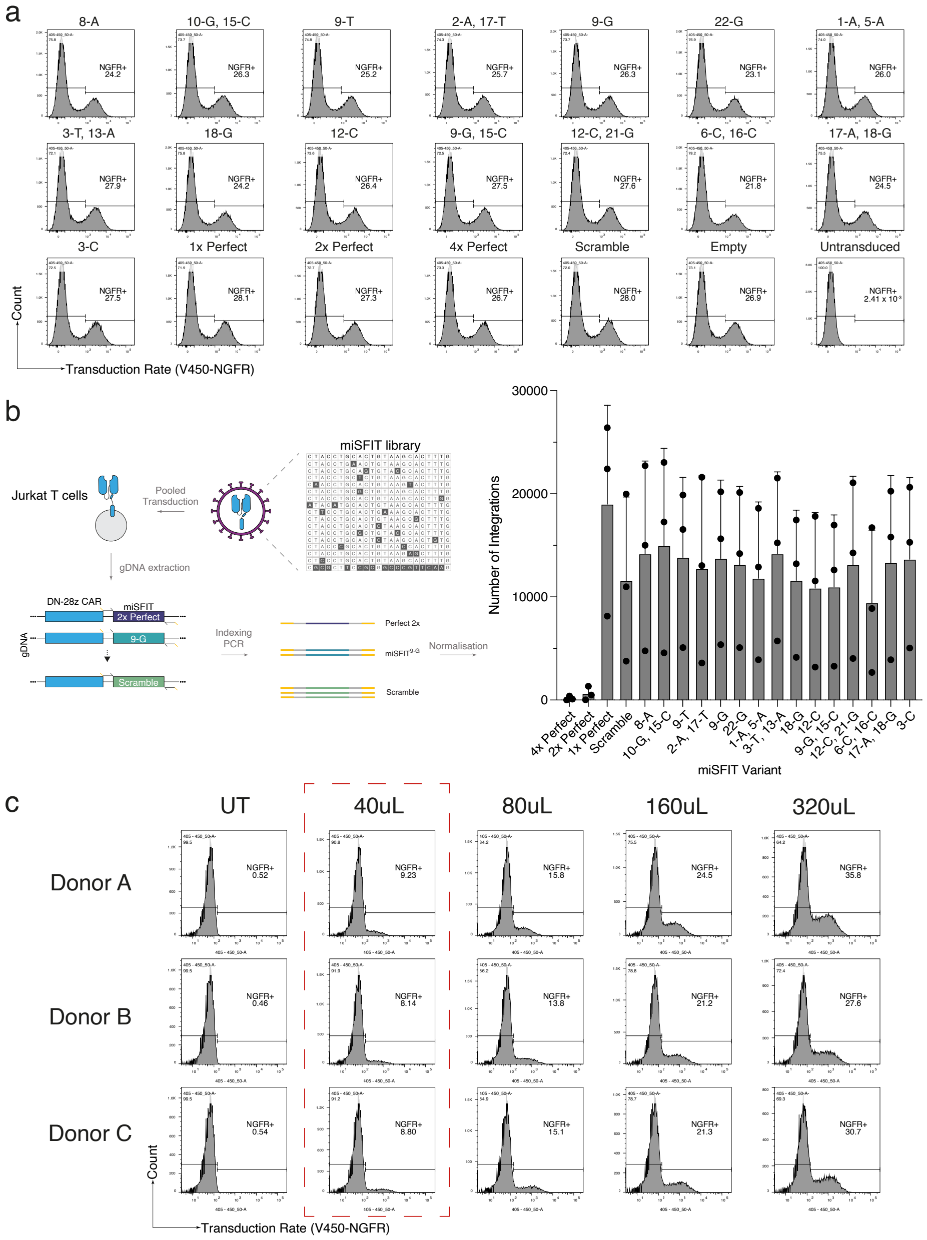


**Extended Data Figure 10**: Creating a lentiviral pool of DN-28z CAR miSFIT for single integrands. To create single cell RNA-sequencing libraries that captured a high dynamic range of DN-28z CAR expression, we created an optimised lentiviral pool that would give rise to a cell population in which each DN-28z CAR miSFIT is equally represented. **a)** Flow cytometry plots of transduction rate, measured by %+ NGFR, used to calculate the functional titre for each DN-28z CAR miSFIT lentivirus. The functional titre was used to calculate the volume of lentivirus to pool. **b)** High throughput sequencing for DN-28z CAR transgene post pooled transduction to validate equal representation for each DN-28z CAR miSFIT. Average integrations by miSFIT. N=3 biological replicates, error bars depict standard deviation. **c)** Titrating volume used to transduce primary T cells with pooled lentivirus for multiplicity of infection $< 0.1$. Flow cytometry plots of transduction rate, measured by NGFR.

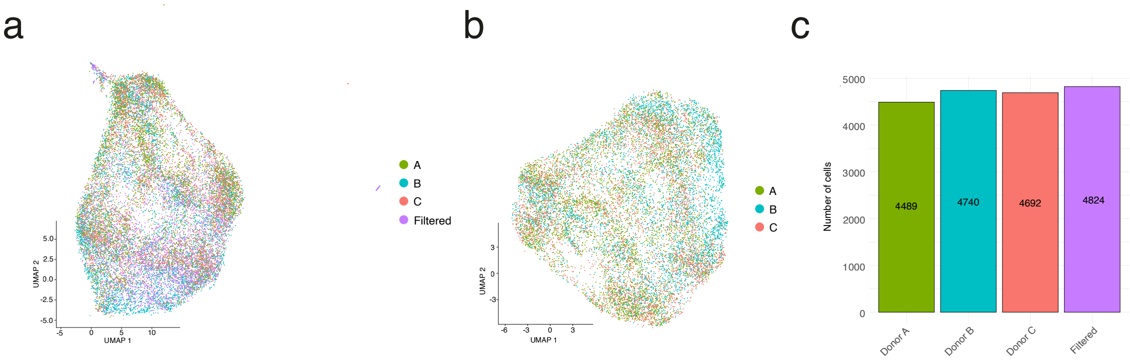


**Extended Data Figure 11**: Genetic demultiplexing reveals homogeneity in transcriptomic response amongst T cell donors in single cell RNA-sequencing. We transduced CD8+ T cells isolated from three human donors and used genetic demultiplexing (**Methods and Materials**) to annotate donor origin for each cell in our single cell RNA-sequencing atlas. UMAP visualisations of genetic demultiplexing results **a)** pre-filtering and **b)** post-filtering. Cells filtered due to being identified as doublet, multiplet, or empty droplet. **c)** Number of cells per donor or filtered.

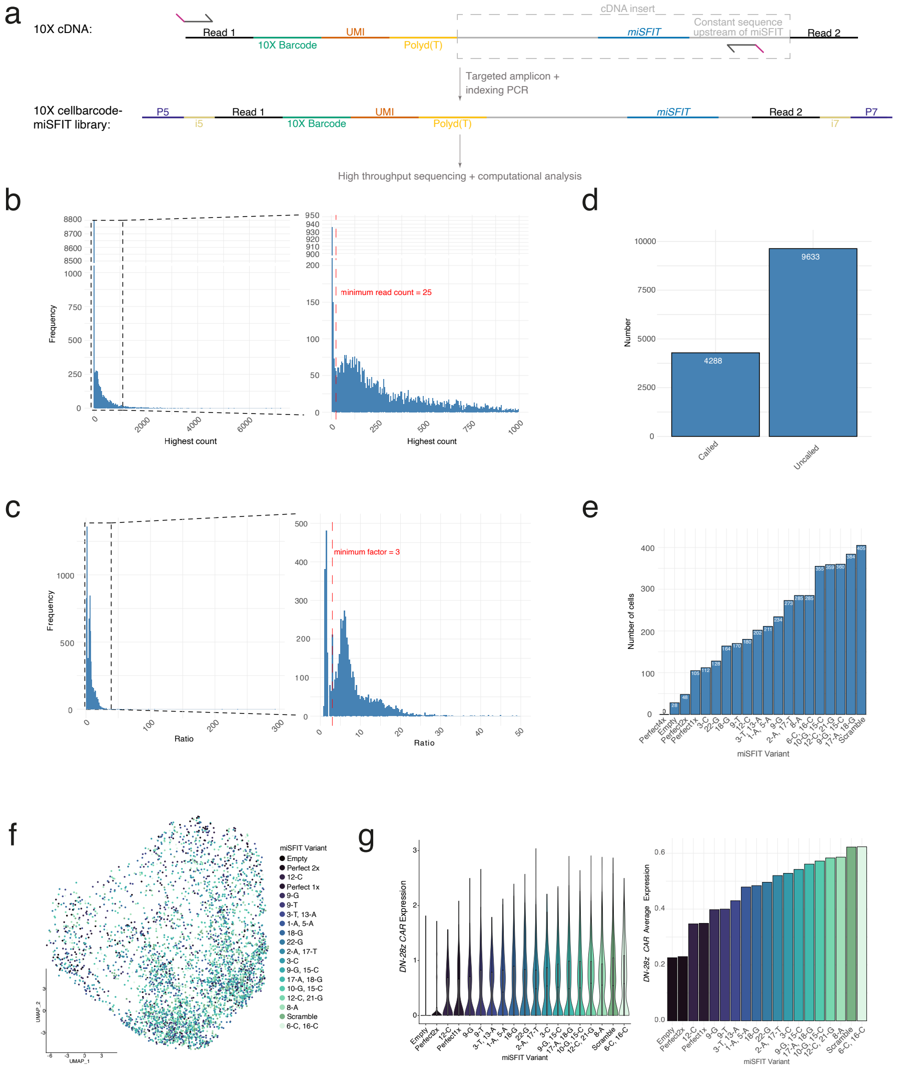


**Extended Data Figure 12**: miSFIT tune CAR expression in single cell RNA-sequencing. **a)** To annotate the cell barcodes of our single cell RNA-sequencing experiment with miSFIT sequence, we used targeted PCR to create a sequencing library that captured the miSFIT sequence and 10X cell barcode. We subjected this library to high throughput sequencing, where Read 1 captured the 10X cell barcode and Read 2 captured the miSFIT sequence. After, we applied a two-step computational algorithm. In the first step, the algorithm conducted the following: (i) filtered the 10x cell barcodes for those that passed quality control (e.g., filtering out multiplets, empty droplets, high mitochondrial counts) and (ii) tallied the number of times each miSFIT sequence appeared for each 10X cell barcode. In the second step, the algorithm then considered the two miSFIT of the highest counts (respectively called M1 and M2) and assigned miSFIT to a 10X cell barcode should the following conditions be met: (1) M1 $\neq$ M2, (2, **(b)**) minimum read count $>$ 25, and (3, **(c)**) M1:M2 ratio $>$ 3. If any of these conditions were not met, the 10X cell barcode would not be assigned a miSFIT sequence. **a-e)** miSFIT demultiplexing. **a)** Schematic of PCR strategy to capture 10X cellbarcode and miSFIT sequence. **b)** Distribution of cells passing minimum read count ($>$25) **(c)** and minimum factor ($>$3). **c)** thresholds for miSFIT demultiplexing (**Materials and Methods**). **d)** Total number of cells demultiplexed and **e)** number of cells demultiplexed per miSFIT. **f)** UMAP visualization of cells demultiplexed by miSFIT. **g)** *DN-28z CAR* distribution and average expression by miSFIT.

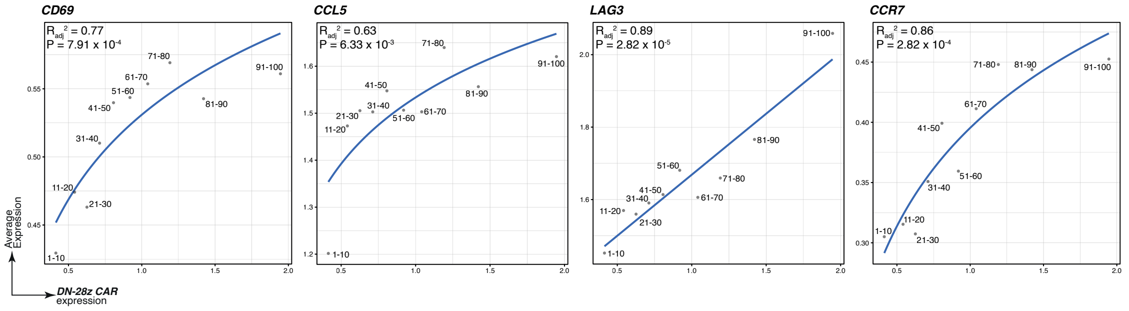


**Extended Data Figure 13**: Cell markers correlate with DN-28z CAR expression. Regression analysis of average expression between *DN-28z CAR* and T cell markers in activation, (*CD69*); effector function, (*CCL5*); exhaustion, (*LAG3*); and memory, (*CCR7*). All genes were fit with a logarithmic model, except *LAG3* which was fit with a linear model. $R_{adj}{}^2$ and P refer to regression analysis model parameters.

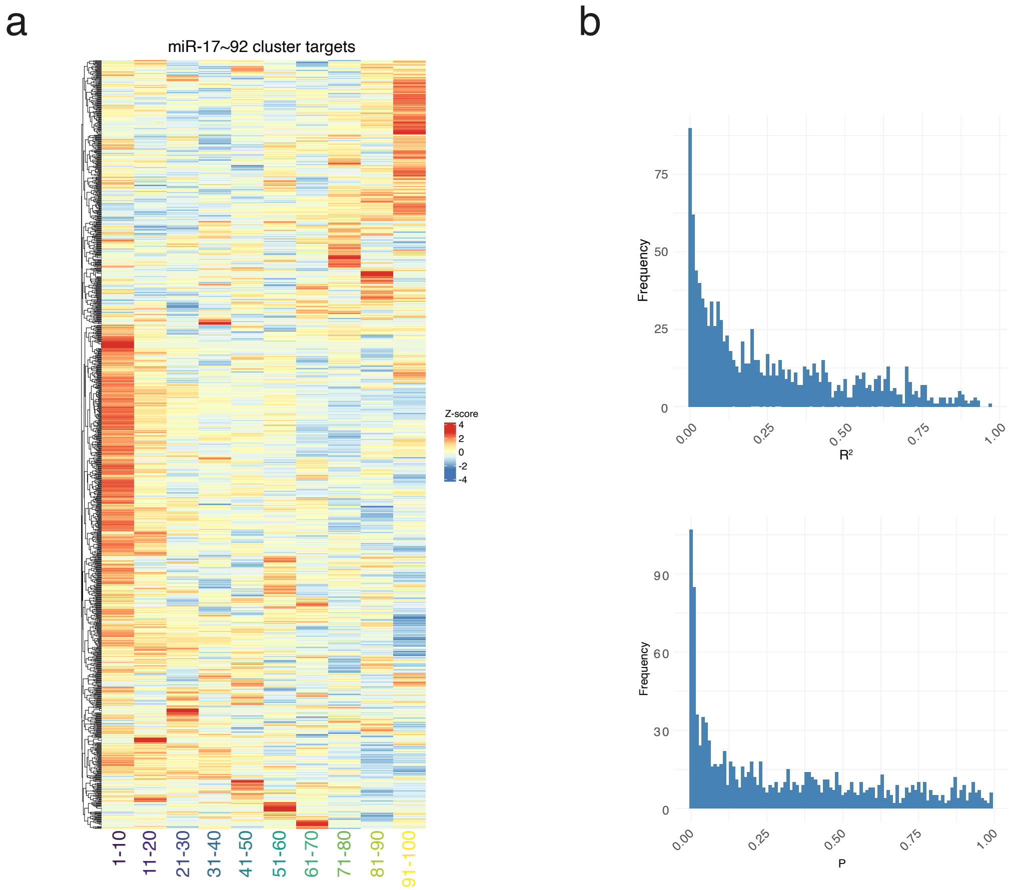


**Extended Data Figure 14**: miSFIT do not impact the miR-17∼92 cluster regulome. To investigate if CAR miSFIT constructs acted as a miRNA sponge, we examined the expression patterns of endogenous miR-17∼92, as predicted by TargetScan (**Supplementary Table 10**, [? ]). **a)** Z-score heatmap and hierarchical clustering of genes predicted to be targeted by miR-17∼92 cluster by TargetScan by CAR expression bin. **b)** Linear regression analysis between *DN-28z CAR* and genes targeted by miR-17∼92 cluster.

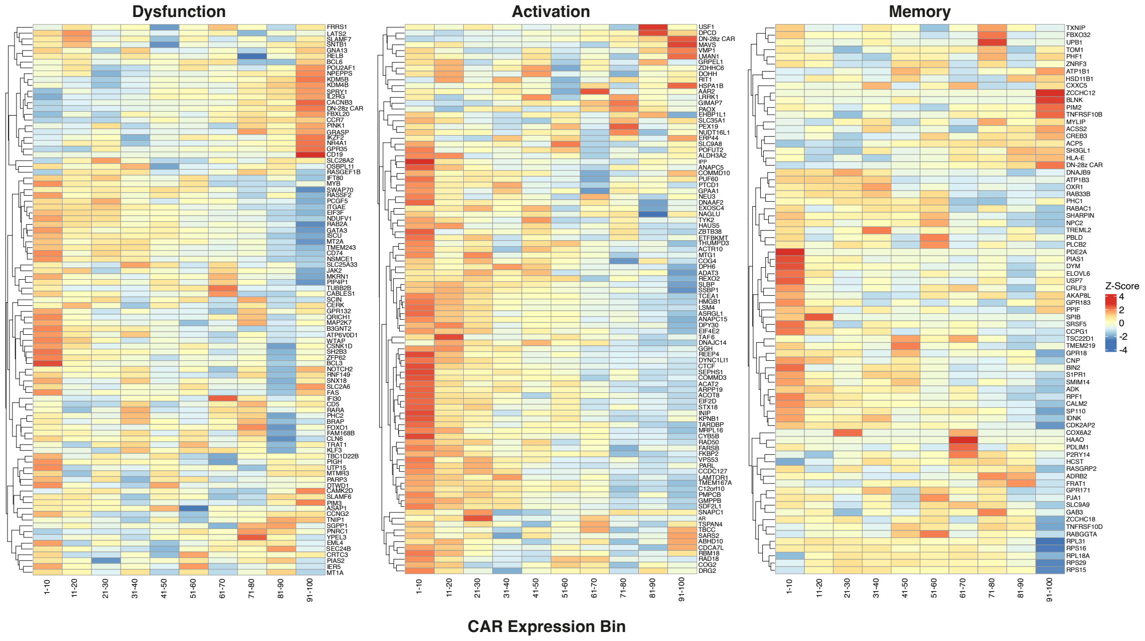


**Extended Data Figure 15**: CAR expression influences modules of activation, dysfunction, and memory phenotypes. To examine CAR expression's impact on transcriptional T cell activation, dysfunction, and memory, we used gene set enrichment analysis (GSEA) on our CAR expression computational bins, using modules published by Singer and colleagues [**?** ]. Z-score heatmap and hierarchical clustering of activation, dysfunction, and memory gene modules. *DN-28z CAR* transcript included in visualisation for reference and was not included when conducting GSEA.

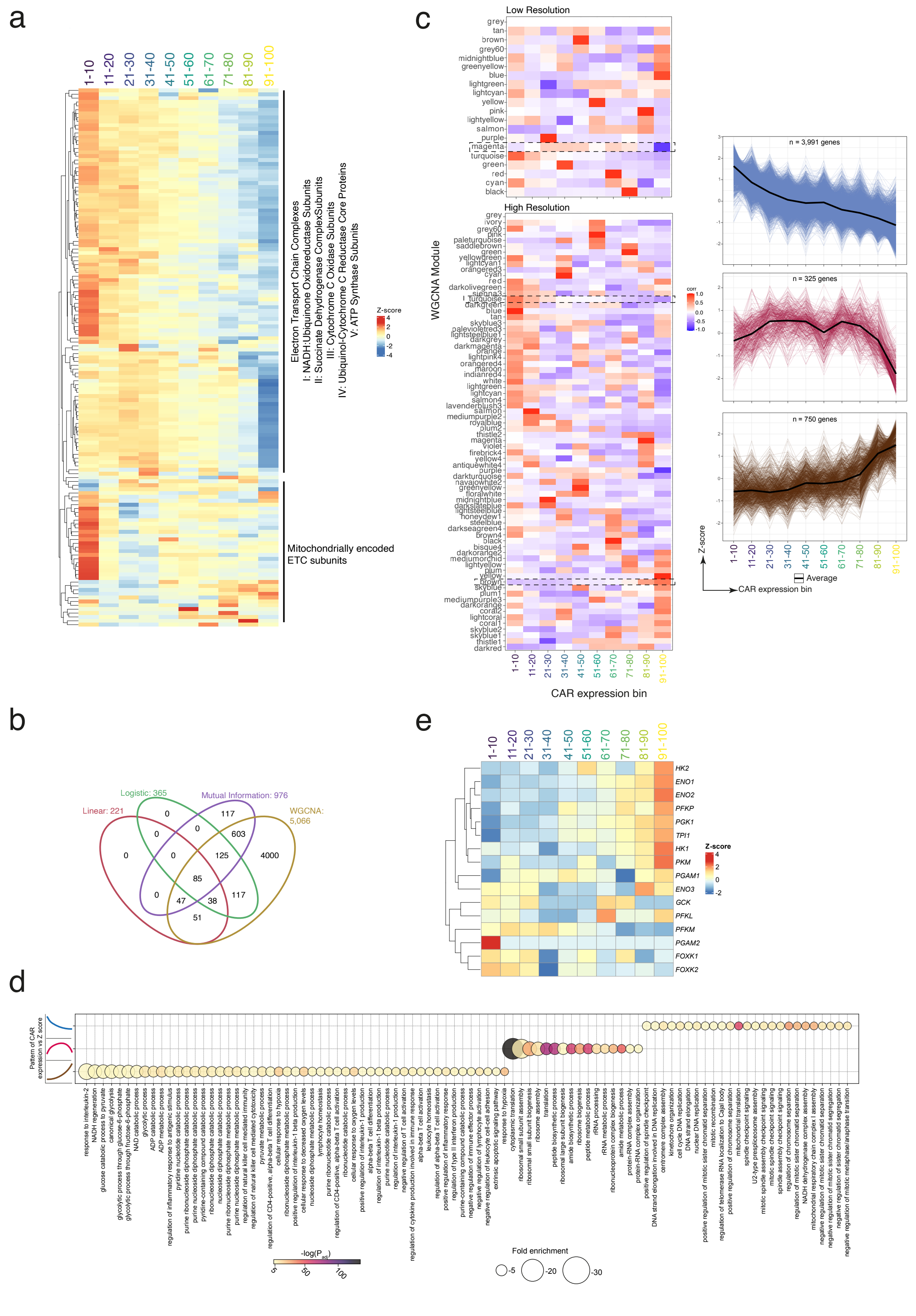


**Extended Data Figure 16**: Excessive CAR expression gives rise to dysfunctional biological processes. **a)** Z-score heatmap and hierarchical clustering of oxidative phosphorylation gene set. Annotated modules of gene families. **b)** Venn diagram of number of genes identified by linear and logarithmic regression, mutual information, and weighted correlation network analysis (WGCNA, [**?**]). **c)** Left: Heatmap of WGCNA modules (named by colour) from high and low resolution analyses. Dotted box indicates investigated modules. Right: Identified three gene expression patterns (top: negative, middle: switch-like, bottom: positive) that correlated with *DN-28z CAR* expression. **d)** Dot plot of gene ontology (GO) themes identified for each WGCNA expression pattern and filtered for a minimum $P_{adj} < 0.5$ (using Bonferroni correction) and fold enrichment > 4. **e)** Z-score heatmap and hierarchical clustering of canonical glycolysis gene ontology gene set.

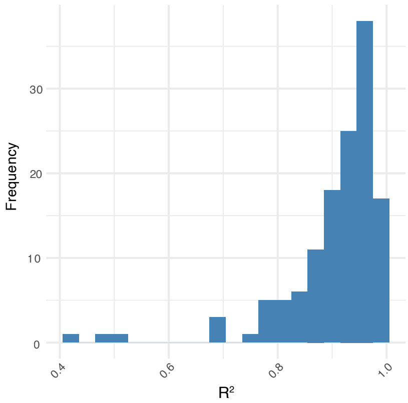


**Extended Data Figure 17**: Regression analysis confirms global, continuous impact on T cell regulators. To determine the relationship between CAR expression and impacted transcription factors, we conducted linear regression analysis between the regulons identified by SCENIC and mutual information and *DN-28z CAR*. $R^2$ refers to goodness of regression fit parameter.
